# Larval microbiota primes the *Drosophila* adult gustatory response

**DOI:** 10.1101/2023.03.14.532561

**Authors:** Martina Montanari, Gérard Manière, Martine Berthelot-Grosjean, Yves Dusabyinema, Benjamin Gillet, Yaël Grosjean, C. Léopold Kurz, Julien Royet

## Abstract

The survival of animals depends, among other things, on their ability to identify threats in their surrounding environment. Senses such as olfaction, vision and taste play an essential role in sampling their living environment, including microorganisms, some of which are potentially pathogenic. This study focuses on the mechanisms of detection of bacteria by the *Drosophila* gustatory system. We demonstrate that the peptidoglycan (PGN) that forms the cell wall of bacteria triggers an immediate feeding aversive response when detected by the gustatory system of adult flies. Although we identify ppk23+ and Gr66a+ gustatory neurons as necessary to transduce fly response to PGN, we demonstrate that they play very different roles in the process. Time-controlled functional inactivation and *in vivo* calcium imaging demonstrate that while ppk23+ neurons are required in the adult flies to directly transduce PGN signal, Gr66a+ neurons must be functional in larvae to allow future adults to become PGN sensitive. Furthermore, the ability of adult flies to respond to bacterial PGN is lost when they hatch from larvae reared under axenic conditions. Recolonization of axenic larvae, but not adults, with a single bacterial species, *Lactobacillus brevis,* is sufficient to restore the ability of adults to respond to PGN. Our data demonstrate that the genetic and environmental characteristics of the larvae are essential to make the future adults competent to respond to certain sensory stimuli such as PGN.

## Introduction

In nature, animals live in a variety of ecological niches colonized by bacteria, viruses and fungi. As a result, animals interact with, and sometimes even host, these co-inhabitants throughout their development and during adulthood. For the better, as these microbial communities can have a positive impact on various physiological parameters of the host such as fertility, longevity and growth, to name but a few ^1–7^. For the worse, as some of these microbes can negatively affect the health and homeostasis of the host ^8^. The ability to detect and respond to these potentially harmful threats is an innate process fundamental to animal survival, that is conserved in all species ^9, 10^. To defend themselves against pathogens, animals have evolved finely tuned cellular and humoral innate immune mechanisms that preserve the physical integrity and health of the host and its offspring ^11, 12^. Defensive responses triggered upon microbial detection come at a cost to the host and might not always be successful ^13, 14 15, 16^. Prior detection of potential dangers in the environment and preparations to face them could be complementary to the canonical responses elicited by pathogens within the host and represent a first line of defense ^17–19^. Thus, the nervous system’s perception of a microbial threat may allow the host to adopt behaviors aimed at reducing the consequences of the infestation on itself and its offspring. These behaviors can act at the individual or population level. Ants and bees have developed social and behavioral immune mechanisms to defend themselves against infection through grooming ^20–22^. Studies in vertebrates and invertebrates have indicated that multiple sensory systems, including olfaction, hearing and vision, are involved in detecting biological threats ^23 18, 19, 24^. In *Drosophila*, the subject of this study, it was found that hygienic grooming behaviors can be induced by fungal molecules or bacterial contact chemicals, via different receptors and neural circuits ^25, 26^. Volatile chemicals, such as geosmin, released by potentially pathogenic fungi can be detected by insects via olfactory receptors and act as repellent, lowering the food intake and modulating the egg-laying rate ^27^. Hallmarks of bacteria are cell wall components such as peptidoglycan (PGN) or lipopolysaccharide (LPS) that are important ligands for receptors allowing eucaryotes to detect and differentiate them from other living organisms ^28^. Interestingly, these receptors are expressed on immune-competent cells as well as on neuronal cells ^18, 29, 30^. Work from several laboratories has shown that bacterial peptidoglycan, an essential component of the bacterial cell wall, mediates many interactions between bacteria and flies ^31–35^. Its recognition by PGRP family members activates NF-κB pathways on immunocompetent cells, leading to the production of immune effectors and regulators. We have recently shown that the same ligand/receptor interactions maintain a molecular dialogue between bacteria and neurons in the central and peripheral nervous systems of flies ^36–38^. We now show that PGN detection by taste neurons triggers an immediate aversive response in adult flies. We identify neurons defined by the expression of the ppk23 gene (ppk23+) as mediators of this bacteria-induced behavior. We also demonstrate that, to be sensitive to PGN, adult flies must hatch from larvae reared in the presence of bacteria and whose Gr66a+ neuronal circuitry is functional. Thus, larval cohabitation with bacteria is a prerequisite for an adult behavioral response to PGN. This demonstrates that the genetic characteristics of the larvae as well as the environmental conditions in which they live are essential to initiate a sensory response to certain molecules later in the adult stage.

## Results

### Bacterial PGN can trigger PER in *Drosophila*

Some Peptidoglycan Recognition Protein family members are expressed in proboscis-hosted taste neurons ^38, 39^. One of them, the membrane receptor PGRP-LC, as well as downstream components of the IMD pathway, are functionally required for transduction of the bacterial PGN signal in these cells ^31, 40, 41^. To test whether this cellular response to a universal bacteria component is associated with a specific fly behavior, we used the proboscis extension response (PER), which is part of the sensorimotor taste circuit in adult flies ^42^. Stimulation of the proboscis by an attractive molecule such as sucrose triggers its immediate and directional extension. When an aversive effect is suspected, the molecule to be tested is combined with a palatable solution such as sucrose. Aversion is measured by the ability of the substance to prevent PER to sucrose. Since our published results ^38^ demonstrated the ability of PGN to stimulate bitter neurons, a cell population involved in aversive behaviors ^43^, we tested whether PGN could suppress the PER response to sucrose. Both Diaminopimelic-type PGN which forms the cell wall of all Gram-negative bacteria and of Bacilli (we used here PGN from *Escherichia coli*, Ec-PGN), and Lysine-type PGN found in the cell wall of Gram-positive bacteria (we used here PGN from *Staphylococcus aureus*, Sa-PGN) were tested ^33^. In PER, Dap-PGN was found to be aversive to reference flies (w-) when used at 100 (PGN100) and 200 µg/mL (PGN200) but not at lower concentrations (Fig 1A). These effects were specific to DAP-type PGN as they were not observed with Lys-type PGN (Fig 1A). Similar aversive effects of DAP-type PGN were obtained with two other commonly used reference *Drosophila* strains, yw and canton-S (Fig 1B and C), and with flies reared on either sugar or AA-rich medium (Fig S1A and S1B).

**Fig. 1.**
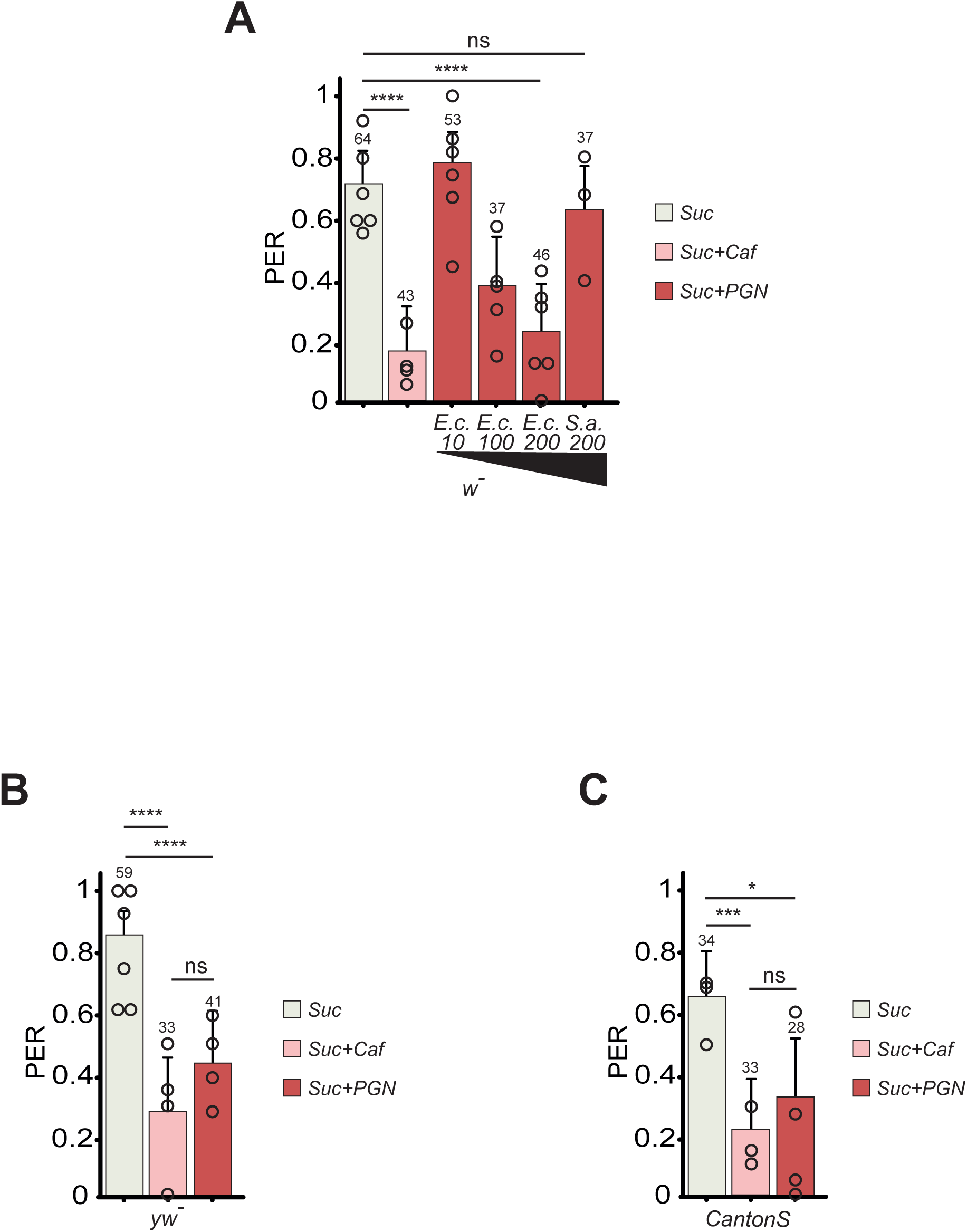
Dose-dependent PGN inhibition of PER. (**A**) PER index of w- flies to control solutions of sucrose 1mM and sucrose 1mM + caffeine 10mM and to sucrose 1mM + increasing concentrations of PGN from *E. coli* K12 (Ec) and *S. aureus* (SA). The numbers below the x-axis correspond to the PGN final concentrations in µg/mL. (**B** and **C**) Aversion to PGN is independent of genetic background. PER index of yw- (**B**) and CantonS (**C**) flies to control solutions of sucrose 1mM and sucrose 1mM + caffeine 10mM and to sucrose 1mM + PGN from E. coli K12 at 200µg/mL. The PER index is calculated as the percentage of flies tested that responded with a PER to the stimulation ± 95% confidence interval (CI). A PER value of 1 means that 100% of the tested flies extended their proboscis following contact with the mixture, a value of 0,2 means that 20% of the animals extended their proboscis. The number of tested flies (n) is indicated on top of each bar. ns indicates p>0.05, * indicates p<0.05, ** indicates p<0.01, *** indicates p<0.001, **** indicates p<0.0001 Fisher Exact Test. Further details can be found in the detailed lines, conditions and, statistics for the figure section.

As with the well-characterized bitter agent caffeine, the ability of PGN to prevent PER was overcome when PGN was mixed with solutions of higher sucrose concentrations (Fig S1C). As the fly legs are taste organs as well ^44^, and PER can also be triggered by chemicals coming into contact with tarsi, we tested PGN in this assay. No aversive response to PGN could be detected in leg-triggered PER, either because the tarsal taste neurons are not involved in the detection of this microbial compound, or because the concentration of sucrose required to trigger the PER on legs overcomes the aversion to PGN (Fig S2A and S2B). We, thereafter, focused on PER triggered by direct contact with proboscis. Because taste neurons have been shown to respond to acidic solutions ^45 46^, we measured the pH of our different PGNs and they were all neutral. This rules out the possibility that the PER response to PGN is due to its acidity (Fig S2C). Taken together, these data demonstrate that Dap-type PGN is perceived as repulsive while Lys-type PGN is not.

### PGN-triggered PER aversion involves Gr66a+ and ppk23+ neurons

To identify the type of neurons that respond to PGN, PERs were performed in flies in which specific groups of neurons were inactivated by overexpression of the inward rectifier potassium channel Kir2.1. To test whether the bitter network was involved in PER aversion toward Dap-type PGN, neurons expressing the pan-bitter taste receptor Gr66a were inactivated ^47^. The ability of Dap-PGN to suppress PER when mixed with sucrose was completely abolished in Gr66a^Gal4^/UAS-Kir2.1 flies demonstrating that Gr66a+ cells are necessary to transduce PGN signal (Fig 2A). Recent work has demonstrated the existence of Gr66a+/ppk23+ and Gr66a+/ppk23- neuron sub-populations that could co-exist or be housed in different sensilla ^48, 49^. These two populations display distinct properties related to salt perception. To assay whether only a subpopulation of neuronal Gr66a+ cells was involved, we used the ppk23^Gal4^ driver. Ppk23^Gal4^/UAS-Kir2.1 flies no longer perceived PGN as aversive demonstrating that Gr66a+ and ppk23+ neurons are both necessary to mediate PGN signal (Fig 2B).

**Fig. 2.**
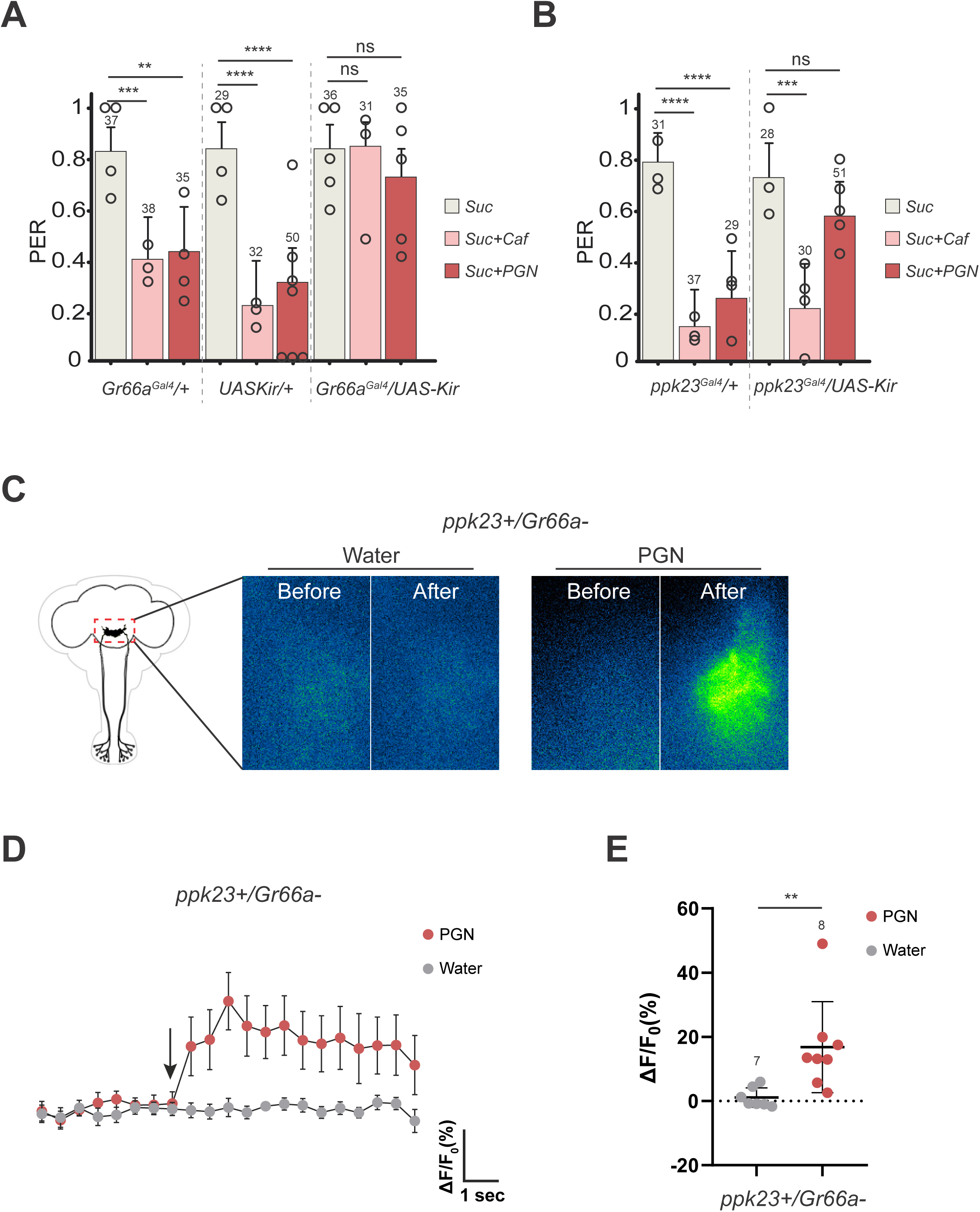
Impairing the activity of Gr66a+ (**A**) or ppk23+ (**B**) neurons via UAS-Kir2.1 abrogates the aversion to PGN. PER index of flies to control solutions of sucrose 1mM and sucrose 1mM + caffeine 10mM and to sucrose 1mM + PGN from *E. coli* K12 at 200µg/mL. (**C-E**) Gr66a-/ppk23+ neurons respond to stimulation with PGN on the labellum. Real-time calcium imaging using the calcium indicator GCaMP6s to reflect the *in vivo* neuronal activity of Gr66a-/ppk23+ neurons (Gr66a^LexA^;LexAopGal80;ppk23^Gal4^/UAS-GCaMP6s) in adult brains of flies whose proboscis has been stimulated with PGN. The expression of LexAop-Gal80 antagonizes the activity of Gal4, thus preventing the expression of GCaMP6S in Gr66a+/ppk23+ neurons. (**C**) Representative images showing the GCaMP6s intensity before and after addition of either the control water or the peptidoglycan. (**D**) Averaged ± SEM time course of the GCaMP6s intensity variations (ΔF/F0 %) for Gr66a-/ppk23+ neurons. The addition of water (n=7 flies) or peptidoglycan at 200µg/mL (n=8 flies) at a specific time is indicated by the arrow. (**E**) Averaged fluorescence intensity of negative peaks ± SEM for control (n=7) and peptidoglycan-treated flies (n=8). For (**A**) and (**B**), PER index is calculated as the percentage of flies tested that responded with a PER to the stimulation ± 95% CI. The number of tested flies (n) is indicated on top of each bar. For each condition, at least 3 groups with a minimum of 10 flies per group were used. ns indicates p>0.05, * indicates p<0.05, ** indicates p<0.01, *** indicates p<0.001, **** indicates p<0.0001 Fisher Exact Test. In (**E**) ** indicates p<0.01 non-parametric t-test, Mann-Whitney test. Further details can be found in the detailed lines, conditions and, statistics for the figure section.

### Adult ppk23+/Gr66a- cells respond to PGN

To directly demonstrate that PGN can activate ppk23+ neurons in the adult, we monitored GCaMP signal in the sub esophageal zone (SEZ) of the brain of females following proboscis exposure to PGN. This brain area processes gustatory input from gustatory neurons located in the proboscis. Since adult Gr66a+ neurons can respond to PGN ^38^ and that some labellar Gr66a+ cells are also ppk23+ ^48^, we used an intersectional genetic strategy to quantify calcium activity modifications only in ppk23+/Gr66a- cells ^48^, a neuronal subpopulation sufficient to mediate the PGN-induced PER suppression (Fig S8B). By quantifying live *in vivo* GCamP fluorescence in adult brains of flies whose proboscis has been stimulated with PGN, we confirmed that ppk23+/Gr66a- cells can directly respond to PGN (Fig 2C-2E). Importantly, these cells did not respond to caffeine but did to high salt, a signature expected for ppk23+/Gr66a- cells.

### Temporal requirement of gustatory neurons for PGN detection during the fly’s lifetime

The putative involvement of several neuronal subgroups in PGN detection prompted us to test whether these different neurons are all required at the same time during the fly’s life. Indeed, although most studies aimed at dissecting how flies perceive their environment via the taste apparatus are performed at the adult stage, including this one, the bitter neurons exemplified by the Gr66a+ population are present throughout fly development ^50, 51^. To determine when the neurons necessary for the PGN-induced PER suppression are required, we took advantage of the Gal4/Gal80^ts^ binary system that allowed us to control the inactivation of taste neurons in a spatially and temporally controlled manner (Fig 3A). Surprisingly, when assessing aversion to PGN, adult flies in which Gr66a+ neurons were functionally inactivated only before the adult stage could no longer perceive PGN as aversive (Fig 3B). In the parallel experiment, inactivation of Gr66a+ neurons in adult flies only, had no effect. In contrast, perception by the adult flies of quinine and caffeine required Gr66a+ neurons to be functional in the adult but not in larvae (Fig S3A). This demonstrates a specific role for Gr66a+ neurons in the life history of the animal, with their requirement only during the larval stage allowing the future adult to show aversion towards Dap-PGN. Since both Gr66a+ and ppk23+ neurons are required to respond to PGN, we tested whether ppk23+ cells are the ones at play in the adults. The data presented Fig 3B demonstrated that, unlike GR66a+ neurons, ppk23+ neurons are functionally required in the adult, but not during larval life for PGN-triggered aversion. These results reveal an unexpected role for larval Gr66a+ neurons in priming the adult response to PGN. For this adult sensory behavior, ppk23+/Gr66a- neurons are sufficient.

**Fig. 3.**
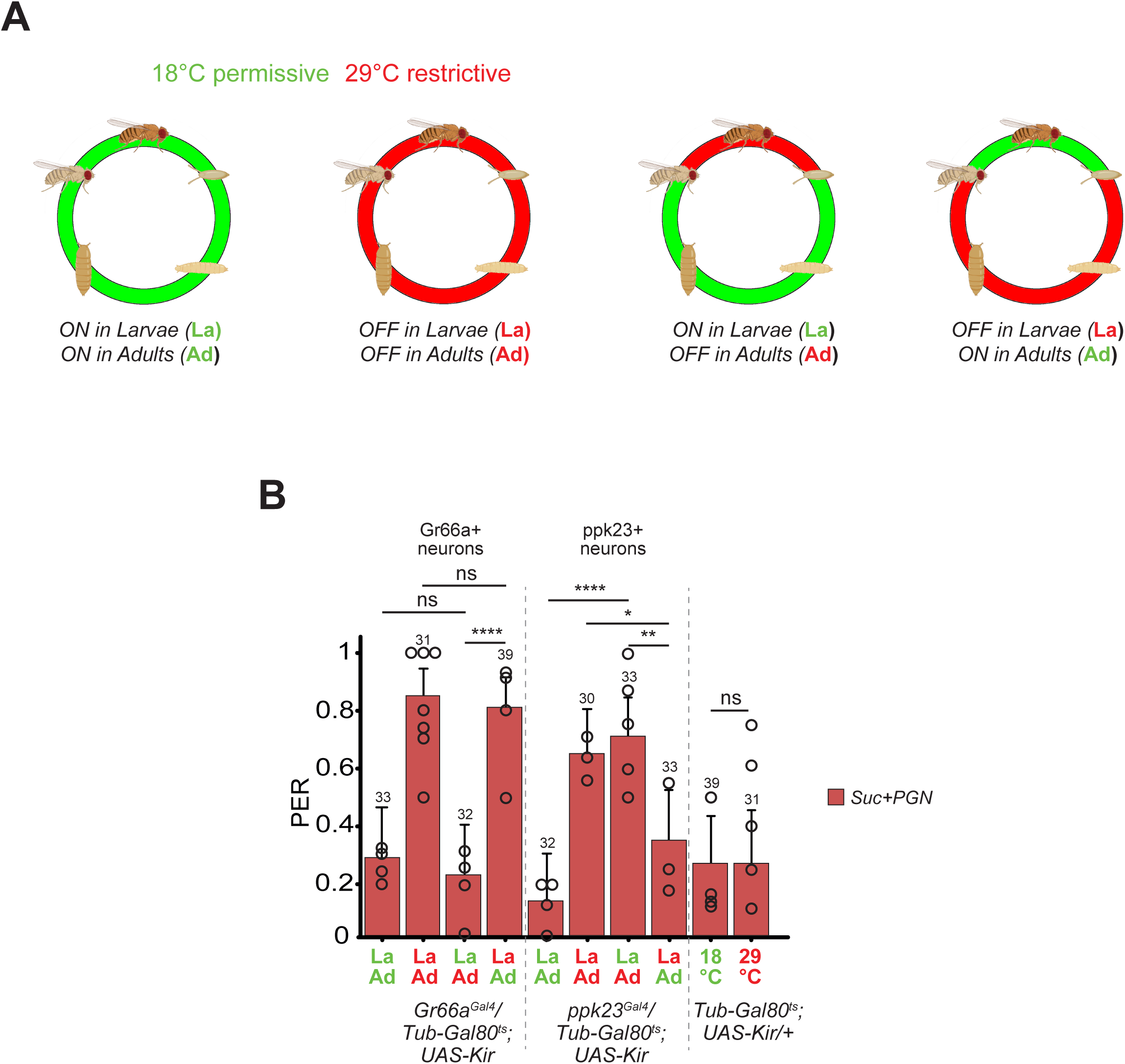
(**A**) Graphical representation of the life periods during which flies are shifted from 18**°**C (green) to 29**°**C (blue). (**B**) Gr66a+ neurons are functionally required in the larval stage for PGN-triggered aversion, but are dispensable during adult life. In opposition to ppk23+ neurons which are required in the adult but not during the larval stages. PER index of flies to sucrose 1mM + PGN from *E. coli* K12 at 200µg/mL. The ubiquitously expressed Tub-Gal80^ts^, that inhibits the activity of Gal4, is temperature sensitive: it’s active at 18**°**C and inactivated at 29**°**C, allowing the expression of UASKir2.1 and the consequent impairment of Gr66a+ or ppk23+ neurons activity. For (**B**), PER index is calculated as the percentage of flies tested that responded with a PER to the stimulation ± 95% CI. The number of tested flies (n) is indicated on top of each bar. ns indicates p>0.05, * indicates p<0.05, ** indicates p<0.01, *** indicates p<0.001, **** indicates p<0.0001 Fisher Exact Test. Further details can be found in the detailed lines, conditions and, statistics for the figure section.

### TRPA1 is functionally required in larval GR66a+ neurons for a response of adult flies to PGN

Using calcium imaging as a readout, we have shown in a previous study that certain components of the IMD pathway (Fig 4A), including the upstream transmembrane receptor PGRP-LC, are functionally required for the transduction of Dap-PGN signal in adult Gr66a+ neurons^38^. We therefore, tested whether this well-characterized PGN receptor was also required in ppk23+ cells. PER responses to Dap-PGN were not affected by RNAi-mediated inactivation of PGRP-LC in either ppk23+ or Gr66a+ neurons (Fig 4B). These results suggesting that Dap-PGN triggered PER is independent of the IMD pathway were confirmed using RNAi against other components of the pathway (Fig S4A and S4B). Consequently, the IMD pathway is neither required in the Gr66a+ larval neurons nor in the adult ppk23+ cells for the PGN-mediated PER. The Gr66a receptor was also tested for its involvement in PGN-induced PER suppression as it is considered, together with Gr33a, as the main co-receptor for the detection of bitter molecules. While Gr66a mutants lost PER aversion for caffeine, they did not for PGN (Fig S4C). In addition to bitter taste receptors, the transient receptor potential TRPA1 channel has been implicated in aversive taste stimulation ^52 53^ . Its RNAi-mediated inactivation in larval Gr66a+ neurons, but not in ppk23+ neurons, abolished the ability of adult flies to respond to PGN (Fig 4C and 4D). These results demonstrate that the expression of TRPA1 receptor in larval Gr66a+ neurons is required for the adults that hatch from these larvae to respond to PGN.

**Fig. 4.**
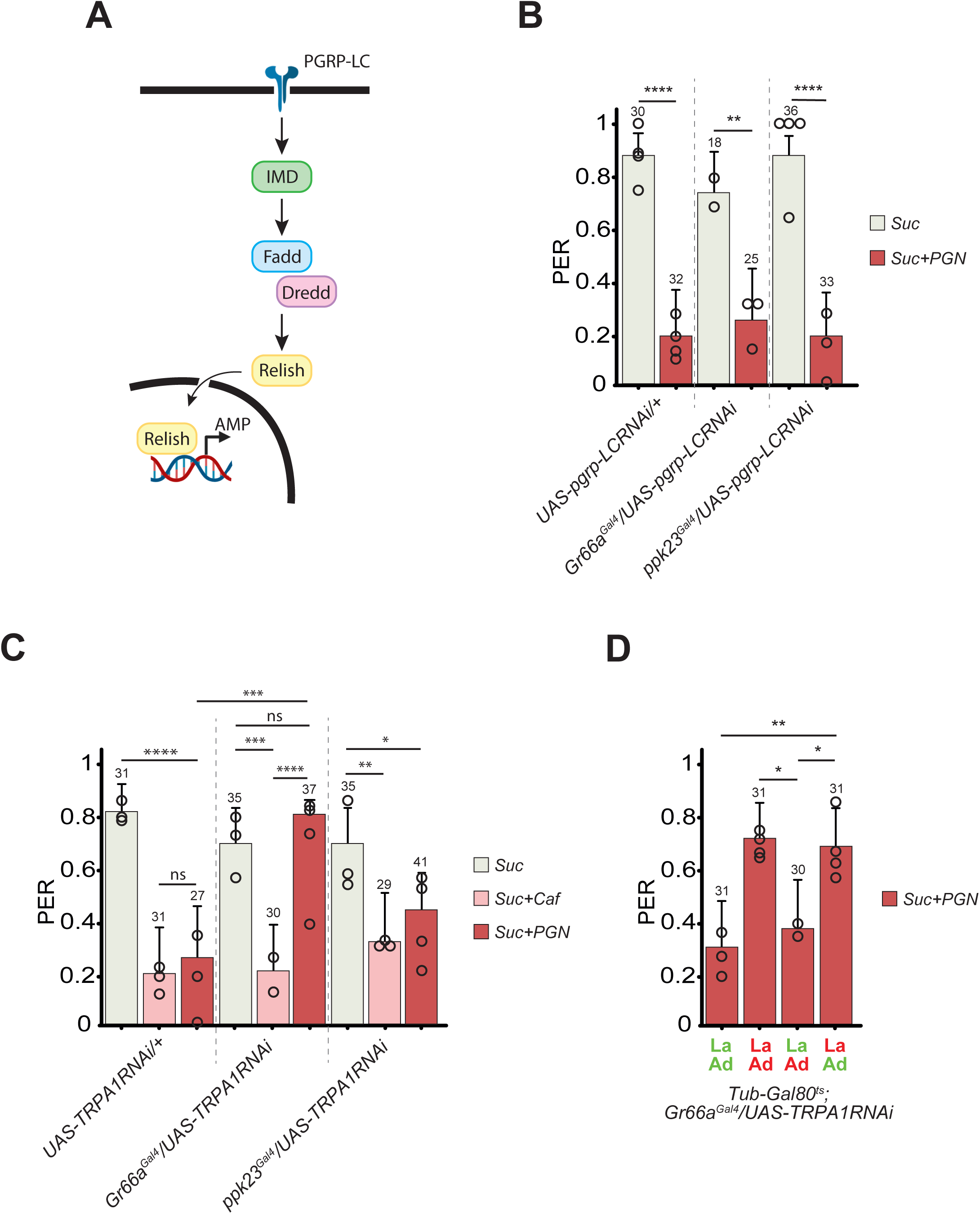
(**A**) Graphical representation of the IMD pathway. (**B**) RNAi-mediated PGRP-LC (UAS-pgrp-LC RNAi) inactivation in the Gr66a+ or ppk23+ cells does not affect PGN-triggered aversion. (**C**) RNAi-mediated TRPA1 (UAS-TRPA1 RNAi) inactivation in the Gr66a+ cells abrogates PGN-induced aversion, while its inactivation in ppk23+ cells has no effect on the aversion phenotype. (**D**) The nociceptive receptor TRPA1 is required in the larval stages for PGN-induced aversion, while its RNAi dependent inactivation in the adult stage has no effect on PGN-triggered aversion. The ubiquitously expressed Tub-Gal80^ts^, that inhibits the activity of Gal4, is temperature sensitive: it’s active at 18**°**C and inactivated at 29**°**C, allowing the expression of TRPA1-RNAi and the consequent inactivation of TRPA1 in Gr66a+ cells. For (**B**), (**C**) and (**D**) PER index of flies to control solutions of sucrose 1mM and sucrose 1mM + caffeine 10mM and to sucrose 1mM + PGN from *E. coli* K12 at 200µg/mL. PER index is calculated as the percentage of flies tested that responded with a PER to the stimulation ± 95% CI. The number of tested flies (n) is indicated on top of each bar. ns indicates p>0.05, * indicates p<0.05, ** indicates p<0.01, *** indicates p<0.001, **** indicates p<0.0001 Fisher Exact Test. Further details can be found in the detailed lines, conditions and, statistics for the figure section.

### Axenic flies are not able to trigger PER aversion to PGN

The involvement of multiple temporal and spatial inputs for PGN detection led us to try to identify the nature of the larval-sensed trigger(s) that prepare the adult response to PGN. Since larvae burrow and live in non-aseptic environments, and the larval microbiota has a strong impact on host immunity, physiology and behavior, we tested its necessity for the adult fly response to bacterially-derived PGN. When flies were reared on antibiotic (axenic) medium, the number of bacteria by fly was strongly reduced (Fig 5A). Interestingly, while axenic flies were still able to respond to caffeine their response to PGN was completely abolished (Fig 5B). Similar results were obtained using different genetic backgrounds demonstrating the universality of the observed effects (Fig S5A and S5B). The PER suppression in response to PGN was also abolished when flies were made axenic by bleaching the eggs (Fig S5C and S5D) eliminating possible side effects of antibiotics on fly taste. These results demonstrate that while exposure to bacteria is not mandatory for the adult response to caffeine, animals must be exposed to bacteria for the adults to perceive Dap-PGN as aversive. To further confirm the inability of axenic adult flies hatched from germ-free larvae to respond to PGN we used calcium imaging. Whereas calcium levels were increased in ppk23+/Gr66a- neurons following PGN exposure to the proboscis of conventionally reared flies, this was no longer the case for axenic flies (Fig 5C-5E). To ensure that aseptic conditions do not, in a non-specific manner, lower the activity potential of neurons, we quantified the salt response that is dependent upon ppk23+ cells. The salt-related activation of these ppk23+/Gr66a- neurons monitored by calcium-imaging was also not affected by germ-free conditions, confirming the specific effect of aseptic conditions upon PGN sensitivity (Fig S5E). These results combined to our temporal genetic inactivation assays suggest that larval Gr66a+ neurons have to be exposed to bacteria for the adult ppk23+ neurons to respond to PGN. We verified that activation of Gr66a+ adult neurons by PGN is involved in a process different form the PER by testing axenic and non-axenic animals. As shown in Fig S5F, adult Gr66a+ cells were activated by PGN in both aseptic and non-aseptic conditions. Consequently, in the adult proboscis, while Gr66a+ cells can respond to PGN regardless of the fly’s life experience with bacteria, ppk23+/Gr66a- neurons can respond to PGN, but non-aseptic conditions are necessary.

**Fig. 5.**
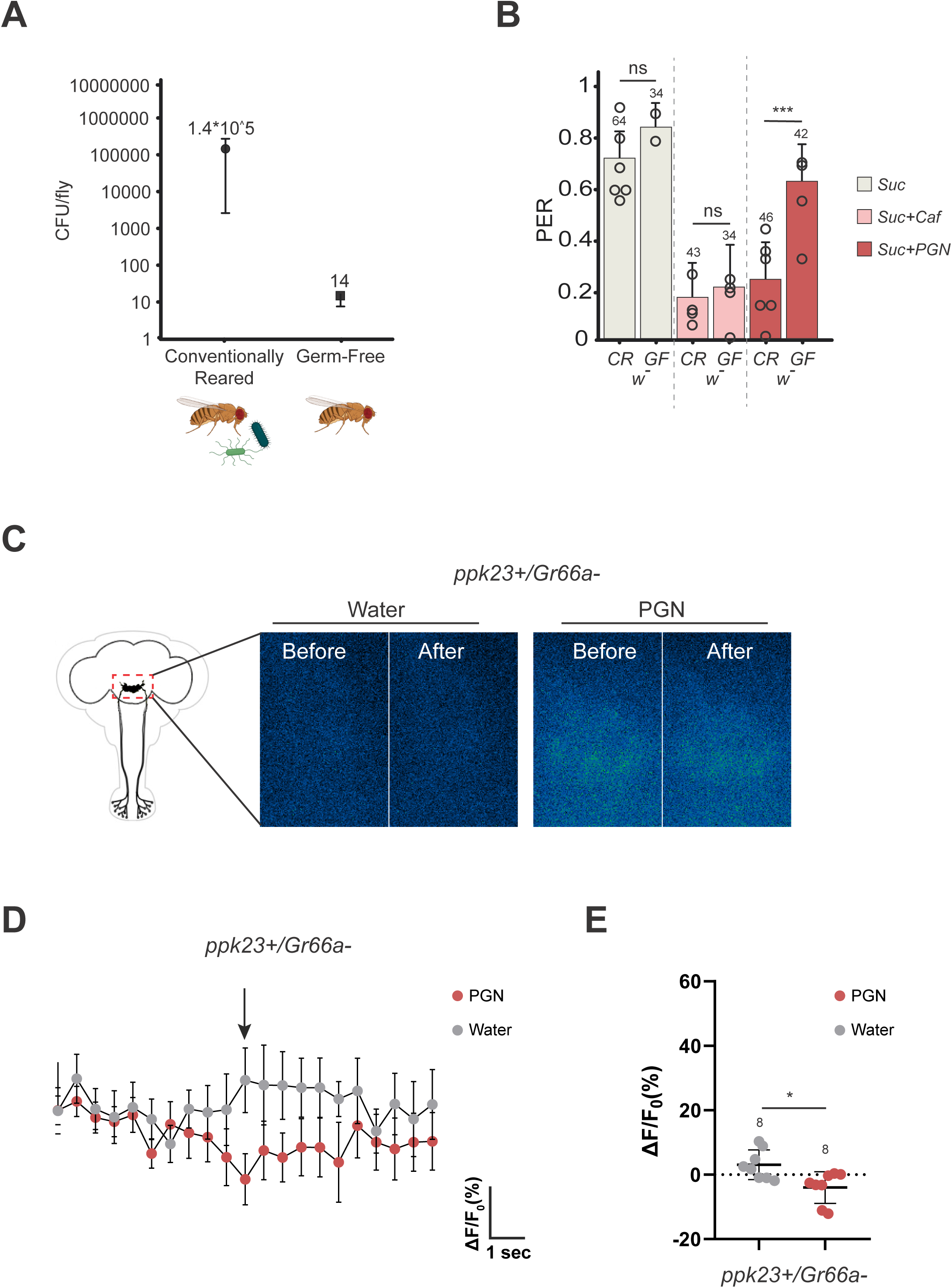
(**A**) CFU/fly comparing conventionally raised flies to those raised on antibiotic media. Antibiotic treatment is effective in reducing the fly bacterial load. (**B**) Germ-free flies do not show PGN-induced PER suppression. PER index of w- Germ-free (GF) and conventionally raised (CR) flies to control solutions of sucrose 1mM and sucrose 1mM + caffeine 10mM and to sucrose 1mM + PGN from *E. coli* K12 at 200µg/mL. The data showed for CR flies are the same as in figure 1A. For (**B**), PER index is calculated as the percentage of flies tested that responded with a PER to the stimulation ± 95% CI. The number of tested flies (n) is indicated on top of each bar. For each condition, at least 3 groups with a minimum of 10 flies per group were used. ns indicates p>0.05, * indicates p<0.05, ** indicates p<0.01, *** indicates p<0.001, **** indicates p<0.0001 Fisher Exact Test. (**C-E**) Real-time calcium imaging using the calcium indicator GCaMP6s to reflect the *in vivo* neuronal activity of Gr66a-/ppk23+ neurons (Gr66a^LexA^; LexAopGal80;ppk23^Gal4^/UASGCaMP6s) in adult brains of flies whose proboscis has been stimulated with PGN. The expression of LexAop-Gal80 antagonizes the activity of Gal4, thus preventing the expression of GCaMP6S in Gr66a+/ppk23+ neurons. (**C**); Representative images showing the GCaMP6s intensity before and after addition of either the control water or the peptidoglycan. (**D**); Averaged ± SEM time course of the GCaMP6s intensity variations (ΔF/F0 %) for Gr66a-/ppk23+ neurons. The addition of water (n= flies) or peptidoglycan (n= flies) at a specific time is indicated by the arrow. (**E**) Averaged fluorescence intensity of positive peaks ± SEM for control (n= 8) and peptidoglycan-treated flies (n= 8) In (**E**), * indicates p<0.05 non-parametric t-test, Mann-Whitney test. Further details can be found in the detailed lines, conditions and, statistics for the figure section.

### Larval microbiota is a pre-requisite to implementing the adult gustatory response to PGN

Because flies are in contact with environmental bacteria throughout development and later in adulthood, we sought to identify the temporal window during which the presence of bacteria impacts the gustatory system’s response to PGN. For that purpose, flies were reared on conventional (non-aseptic) medium and transferred to antibiotic-containing medium at different periods of their life cycle and for different durations (Fig 6A). All emerged flies were then tested for their ability to respond to PGN. Whereas flies emerged from non-aseptic larvae reared immediately after hatching on antibiotic medium responded adversely to PGN (Fig 6B), those from larvae reared on antibiotic (aseptic) medium lost this ability, despite adults’ exposure to non-aseptic environment (Fig 6B). To further reduce the permissive time window, larvae were reared on a non-aseptic medium for only a part of the larval period. The efficacity of the different treatments were monitored using CFU tests (Fig S6A). Exposure of larvae to antibiotic food for the 72 first hours of development was sufficient to abolish the aversive response of the adult to PGN (Fig 6C). Similarly, rearing germ-free eggs on non-aseptic medium for the first 48 hours of development was sufficient to restore PGN responsiveness in adults (Fig 6D). Furthermore, exposure of axenic larval pupae to bacteria (Fig S6B and S6C), as well as exposure of germ-free larvae and adults to PGN-containing media (Fig S7A) was not sufficient for the adult’s PGN-induced PER suppression to be re-established. These results indicate that cohabitation of early larvae with bacteria is a prerequisite for the adult taste aversive response to PGN to be established. In addition, these data are consistent with a role for Gr66a+ taste neurons only during larval life for the adult to respond adversely to PGN in a PER assay.

**Fig. 6.**
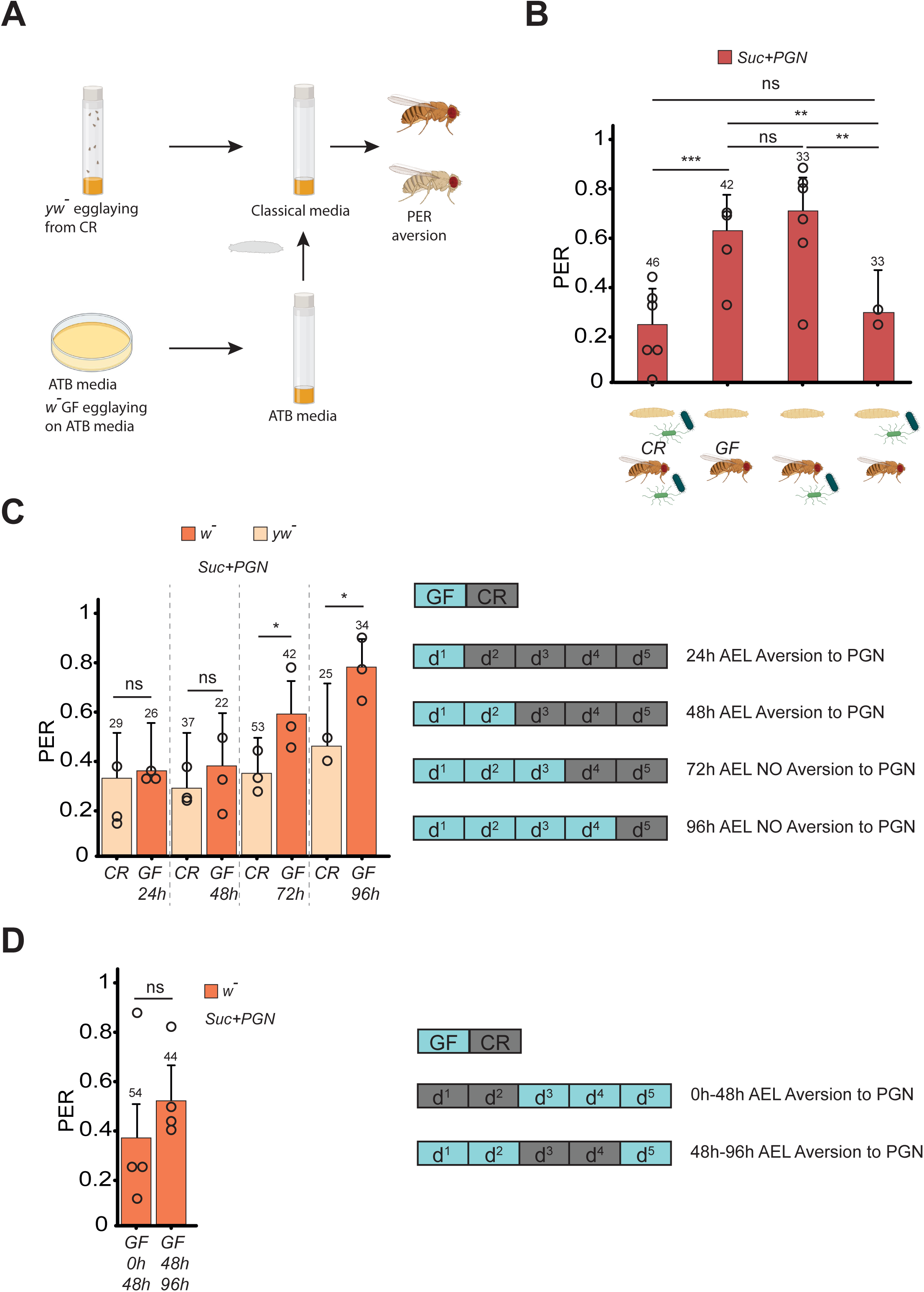
(**A**) Graphical representation of the protocol to contaminate Germ-free larvae at different stage of the larvae development. Two ovipositions of yw^-^ flies on standard medium and w- flies on antibiotic-enriched medium started simultaneously. After 4h the embryos of the w^-^ flies are sterilized using bleach and transferred onto antibiotic-enriched media. Upon reaching the desired stage of development the w^-^ germ-free larvae are transferred to the same tube in which the yw^-^ larvae are growing. Once the adult stage is reached, flies are separated according to body color and their PER response assayed. (**B**) Co-habitation with bacteria only during the larval stage is sufficient to trigger PGN-induced PER suppression. PER index of w- flies, shifted from Germ-free to conventional raising conditions (and the reverse) upon pupation, to control solutions of sucrose 1mM + PGN from *E. coli* K12 at 200µg/mL. The data showed for conventionally raised (CR) and Germ-free (GF) flies are the same as in figure 1A and 5B. (**C**) PER index of w- Germ-free animals, shifted to conventional raising conditions at different stages of their larval development (as in fig. 6A), to solutions of sucrose 1mM + PGN from *E. coli* K12 at 200µg/mL. The graphic represents how many days after the egg laying (AEL) larvae were transferred to yw^-^ used media, and their response to stimulation with PGN. (**D**) PER index of w- Germ-free animals, transferred for short periods of time of their development under conventional raising conditions, to solutions of sucrose 1mM + PGN from *E. coli* K12 at 200µg/mL. The graph represents the timing of the shift from one condition to another. For (**B**), (**C**) and (**D**), PER index is calculated as the percentage of flies tested that responded with a PER to the stimulation ± 95% CI. The number of tested flies (n) is indicated on top of each bar. For each condition, at least 3 groups with a minimum of 10 flies per group were used. ns indicates p>0.05, * indicates p<0.05, ** indicates p<0.01, *** indicates p<0.001, **** indicates p<0.0001 Fisher Exact Test. Further details can be found in the detailed lines, conditions and, statistics for the figure section.

### Mono-association of larvae with *Lactobacillus brevis* is sufficient to restore the aversive adult response to PGN

Although much simpler than that of mammals, previous work has shown that the *Drosophila* microbiota can host about 20 species mainly belonging to the genus *Acetobacter* or *Lactobacillus*, depending of the fly’s diet ^54–58^. To test whether a specific bacterial species could mediate larval priming effects, we sequenced the bacteria present in our laboratory fly colony (Fig 7A). Of the few species identified, *Lactobacillus brevis* was one of the most abundant and caught our attention because it has been shown to affect the behavior of the flies it inhabits ^59, 60^. Strikingly, PGN-induced PER suppression was restored in axenic adults obtained from larvae mono-associated with *Lactobacillus brevis* (Fig 7B and 7C). When the same experiment was performed with a related species, *Lactobacillus plantarum*, no PER inhibition was observed (Fig 7C). It has been previously reported that the presence of *L. brevis* in adult animals can alter the fly behavior by modulating the production of octopamine, a neurotransmitter involved in oviposition, male fighting and locomotor activity ^60 61, 62^ . In our case, exposure of adults to *L. brevis* was not sufficient to restore the PGN PER phenotype and we ruled out the putative involvement of octopamine by demonstrating that the PGN is still perceived as aversive by mutants unable to synthetize octopamine (TβH) ^63^ (Fig S7B).

**Fig. 7.**
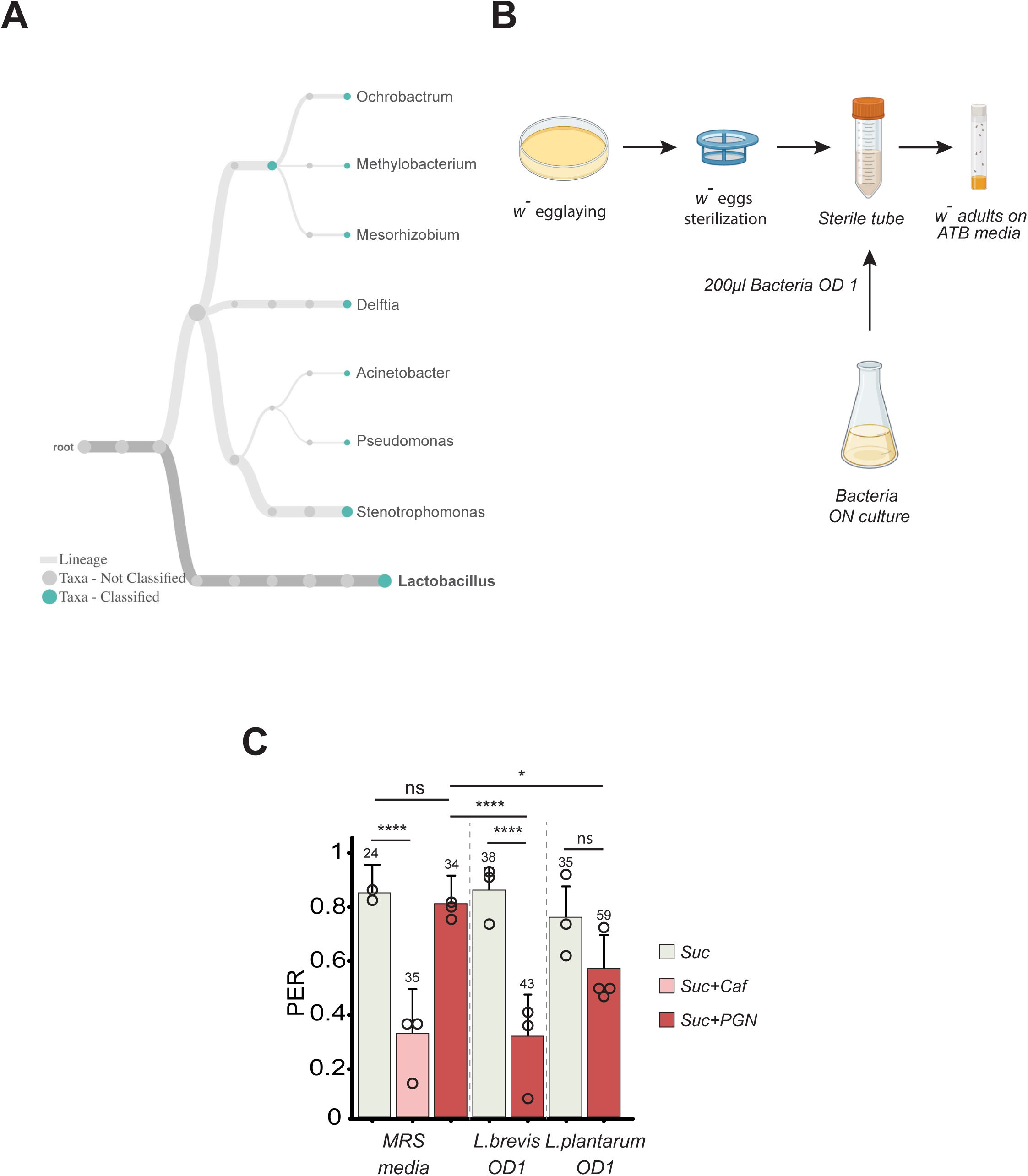
(**A**) Lactobacilli are among the most abundant genera in our conventional raising conditions. Graphical representation of the bacterial genera present in our conventional raising conditions according to 16S sequencing. (**B**) Graphical representation of the protocol for the monoassociation of germ-free flies with a bacteria. After oviposition the embryos are sterilized with bleach and transferred under sterile conditions into a fresh sterile tube then 200µL of bacterial culture at OD 1 is added. Adults emerging from pupae are then transferred on antibiotic-enriched media. (**C**) Mono association of Germ-free flies with *L. brevis*, but not *L. plantarum*, is sufficient to trigger the PGN-induced PER suppression. PER index of w- flies mono-associated with *L. brevis*, *L. plantarum* or exposed to their culture media (MRS) to sucrose 1mM and sucrose 1mM + caffeine 10mM and to sucrose 1mM + PGN from *E. coli* K12 at 200µg/mL. For (**C**) PER index is calculated as the percentage of flies tested that responded with a PER to the stimulation ± 95% CI. The number of tested flies (n) is indicated on top of each bar. For each condition, at least 3 groups with a minimum of 10 flies per group were used. ns indicates p>0.05, * indicates p<0.05, ** indicates p<0.01, *** indicates p<0.001, **** indicates p<0.0001 Fisher Exact Test. Further details can be found in the detailed lines, conditions and, statistics for the figure section.

Overall, these results demonstrate that a period of cohabitation between larvae and specific bacterial species, such as *Lactobacillus brevis*, is mandatory for the emerging adult to perceive PGN as an aversive molecule by ppk23+ neurons.

## Discussion

The data presented here demonstrate that flies are competent to perceive and respond to PGN of bacterial origin via ppk23+ neurons exposed to the environment. However, this adult behavior only takes place if the larvae that will give birth to these flies carry functional Gr66a+ neurons and are reared in the presence of certain bacterial species such as *Lactobacillus brevis*.

### Are Gr66a+ and ppk23+ subpopulations sufficient to mediate larval priming and adult response to PGN?

The sensilla of the proboscis are the main taste detectors and act as straws sampling the outside world. Divided into three classes (S, L, I), they each house several cells, including taste neurons whose dendrites are exposed to the outside world via the sensilla opening ^64^. While Gr66a+/ppk23- neurons are hosted in I- and one subset of S-sensilla (called Sb), Gr66a+/ppk23+ neurons are found in the other subset of S-sensilla (called Sa). Gr66a-/ppk23+ cells are present in few I-sensilla, almost all S- and all L-sensilla ^48^. Therefore, some sensilla host both Gr66a+/ppk23+ and Gr66a-/ppk23+ cells. Since our data show that Gr66a+ neurons are dispensable and ppk23+ cells essential for PGN-mediated PER, we could conclude that Gr66a-/ppk23+ cells are the only cells involved. However, although Gr66a-/ppk23+ subset can respond specifically to PGN (Fig 2C-2E), flies in which Gr66a-/ppk23+ cells are specifically silenced remain able to sense PGN (Fig S8A and S8B). This suggests that two populations are capable of independently triggering the PGN-mediated PER: Gr66a-/ppk23+ and Gr66a+/ppk23+ neurons. The description of new receptors expressed on specific subsets of Gr66a-/ppk23+ neurons such as Ir7c should help to understand the complexity of this taste language ^65^. A recent study related to salt detection illustrates this complexity. While high concentrations of monovalent salt cause aversion via Gr66a+ and Gr66a-/ppk23+ (Gr66a-/Ir7c-/ppk23+) neurons, high concentrations of divalent salt activate only a subset of Gr66a-/Ir7c-/ppk23+ cells^65^. As with adults, the minimal subset of cells required in larvae to prime the adult response to PGN remains partially elusive. Larval Gr66a+ neurons can be divided into Gr32a+ and Gr32a- subpopulations ^66^. Gr32a>Kir2.1 animals respond to PGN like controls, confirming that adult Gr66a+ neurons, which are all Gr32a+ in flies ^67^, are dispensable for PGN aversion to PER. This also suggests the involvement of the larval Gr66a+/Gr32a- subpopulation (Fig S8C). Silencing other smaller neuronal subgroups using Gr Gal4 drivers ^66^, had no impact on PGN aversion in the adult (Fig S8C). Although our data show that ppk23+ cells are not required during the larval period, the existence of larval Gr66a+/ppk23+ neurons has not been clearly established.

### What receptor(s) transduce the PGN signal in adult ppk23+ neurons?

The demonstration that adult ppk23+ neurons respond to PGN prompted us to identify the upstream receptor(s). Despite the expected involvement of the classical immune PGN sensor PGRP-LC, its functional inactivation has no impact on aversion towards the microbial compound (Fig. 4B). Even elimination of central elements of the signaling pathway, such as Fadd, were without consequences. The lack of involvement of the IMD/NF-kB pathway in the larval priming phase was also demonstrated for larval Gr66a+ neurons. Our data rather identify TRPA1 as a putative receptor necessary on larval Gr66a+ neurons to sense a ligand related to microbial activity. As for the adult response, the yet to be identified receptor is expected to be expressed on the dendrites of adult ppk23+ neurons, exposed to the sensilla-bathing medium and hence to PGN coming from exogenous bacteria. Unfortunately, the physiology of ppk23+ neurons is mostly understood via their role in pheromone perception ^68^ and of high salt avoidance ^48, 65^. Few surface receptors have been described and include Gr66a, ppk23 and Ir7c. We have shown that Gr66a is not required. An alternative approach to identify the receptor could be to decipher the signaling pathway required for PGN-mediated PER in ppk23+ cells. For example, if Gαq is involved, GPCRs would appear as good candidates ^69^.

### How do *Lactobacillus brevis* activate larval priming?

Our results highlight how the genetic characteristics of the larva and the environment in which they live can impact the development of a sensory response in the adult to whom they will give birth. For the priming to take place larvae must co-live with bacteria, while their presence during adulthood is dispensable. If recent reports describe the influence that microbiota has on certain adult fly behaviors ^60, 62^, none of them report on the bacterial requirement specifically in larvae. In many studies associating behavioral changes to septic conditions, the neurotransmitter octopamine (OA) was shown to play a central role. This does not seem to be the case here since *Tϕ3H* mutant flies, which do not produce OA, respond well to PGN. Our data point to changes initiated by the interaction between TRPA1 receptor expressed on larval Gr66a+ neurons and bacteria (at least *Lactobacillus brevis*) present during the first 2-days of the larval life. An important question to address is how such an interaction is taking place. Among many possibilities, one can propose, i) modifications of the media by the bacteria, ii) changes in the gut physiology upon bacteria detection, iii) impact of microbial activity on the host iiii) direct effects of some bacterial components. In the latter category, one candidate is the PGN itself which plays key role in the interactions between flies and bacteria. However, although PGN is universally present on all bacteria, only some of them are able to mediate priming as mono association with *L. plantarum* was not sufficient. Furthermore, our unsuccessful attempts to rescue adult response by incubating larvae with purified PGN do not support this hypothesis. Intriguingly, while the strain of *Lactobacillus plantarum* that we used, considered as a symbiont, was not able to recapitulate the effect of a septic media, Lactobacillus *brevis* that is a pathobiont with a distinctive molecular signature involving uracil was sufficient to make competent adults ^70, 71^. Further work will be necessary to molecularly dissect this interaction.

### What is missing in adult ppk23+ neurons hatch form axenic larvae?

Our data suggest that some pieces of information acquired during the larval life experience are transmitted to the adult. Interestingly this effect shows some degree of specificity. Indeed, while the ppk23+ cell-dependent aversion to PGN is lost, adult ppk23+ neurons from axenic larvae are not lost or anergic as they remain responsive to salt. In addition, the activation of neurons directly exposed to the environment and the lack of this activity when the larvae were reared in axenic conditions suggests that the competence and the changes occur directly within the ppk23+ neurons and not indirectly via other neurons. However, the situation might be more complex as the *in vivo* imaging is performed in the brain where multiple axons converge making the resolution at the neuron-scale difficult. Indeed, a subset of Gr66a-/ppk23+ cells at the periphery might be dedicated to PGN-sensing and not especially to salt, and fully incapacitated by axenic rearing while the others only respond to salt, a capacity unaffected by microbial exposure.

### How can the larval experience influence an adult behavior?

This study raises the question of the mechanisms by which information acquired by larvae in contact with bacteria are specifically transmitted to adults. Since larval Gr66a+ neurons are necessary for this process, it is conceivable that they persist in the adult. However, like most taste neurons that are either lost or transformed during metamorphosis, larval Gr66a+ neurons do not persist into adulthood, except for a subset that becomes adult pharyngeal taste cells ^72 73 74 75^. However, since PER is performed by sensory touch without ingestion and adult Gr66a+ cells are dispensable for the phenotype, this is unlikely to be the case. Another possibility would be that some larval Gr66a+ cells become adult ppk23+ neurons. The activation state of the larval GR66a+ neurons will then have a direct impact on that of the adult ppk23+ neurons.

A long-term memory or a memory maintained throughout metamorphosis or pre-imaginal conditioning could be considered. Definitive proofs of such mechanisms are still lacking since chemicals used to condition and train the larvae could remain in trace form in the media or on the resulting pupae and, in term, influence the newborn flies ^76, 77^. These caveats are not valid here since the spatial and temporal genetic inactivation of neurons and the silencing of TRPA1 unambiguously demonstrate the role of these cells specifically and only during larval life. In addition, the effect of time-controlled bacterial exposure and the lack of response to PGN in adults exposed for five days or from pupae to septic media support the idea that larvae must be reared in non-axenic conditions to produce PGN responsive adults. Bacteria-larvae interactions taking place during the priming period could activate some kind of memory. Such hypothesis is not supported by a recent study that tracked the fate of neurons involved in memory finding no anatomical substrate for a memory trace persisting from larva to adults ^75^. Conditions altering memory consolidation in general will be tested in the future ^78^.

Epigenetic modifications such as histone mark alterations could be affected by the presence of bacteria in the larval gut. Previous reports indicate that defects associated with histone demethylase kdm5 inactivation can be partially rescued by modulating the gut microbiota either by supplementation with *L. plantarum* or by antibiotic treatment ^79^. If epigenetic modifications are involved, they will have to be limited since the changes we detected in adult taste are not generic for all tastants. However, such a possibility should not be ruled out as it has already been shown that animal exposure to microbe can influence behavior and vertical transmission of the phenotype via epigenetic modifications ^80, 81^.

### Multiple inputs of bacterially-derived PGN on taste neurons

Using calcium imaging, we previously demonstrated that PGN exposure impacts neuronal activities in an IMD/NF-κB pathway dependent manner ^38^ . Both brain octopaminergic neurons and adult Gr66a+ taste neurons present IMD pathway dependent calcium modulation following PGN exposure. In the current report we demonstrate that Gr66a+ cells are dispensable for the behavior in the adult stage. Different elements support the fact that the proboscis Gr66a+ cells activation we previously described is linked to a phenotype yet to be fully uncovered. First, when the Gr66a+ are silenced specifically during the adult stage, while the animals no longer respond to caffeine, we still observe the PGN-mediated PER. Second, when the animals no longer react to the PGN for they have been raised in axenic conditions, the Gr66a+ neurons still respond to the presence of PGN. Finally, while the IMD pathway was essential for the increased calcium concentration in Gr66a+ neurons following PGN exposure, this signaling cascade is not required for the PGN-mediated PER. In agreement, we’ve identified that another set of neurons present in the proboscis is necessary and sufficient for the PGN-mediated PER, the ppk23+ neurons. Further work will be needed to understand how flies integrate these responses to PGN and to other bacterial components such as lipopolysaccharide to respond adequately to the presence of bacteria in their immediate surrounding.

## Author contributions

M.M., G.M., L.K, Y.G. and J.R. conceived the experiments. M.M., G.M., M.B.G., I.D., G.G., L.K. performed the experiments. M.M., L.K. and J.R wrote the manuscript. Y.G. and J.R. secured funding.

## Competing interests

The authors have declared that no competing interests exist.

## Acknowledgments

We thank François Leulier, Sandrine Hughes and Maxime Veites for their help with microbiota sequencing. We thank Emilie Avazéri and Gladys Gazelle for their technical help. This work was supported by CNRS, ANR BACNEURODRO (ANR-17-CE16-0023-01), Equipe Fondation pour la Recherche Médicale (EQU201603007783) et l’Institut Universitaire de France to J.R. and the ANR Pepneuron (ANR-21-CE16-0027) to J.R. and Y.G. Research in Y.G.’s laboratory is supported by the CNRS, the “Université de Bourgogne Franche-Comté”, the Conseil Régional Bourgogne Franche-Comté (PARI grant), the FEDER (European Funding for Regional Economical Development), and the European Council (ERC starting grant, GliSFCo-311403)

## Figure Legends

**Fig. S1.**
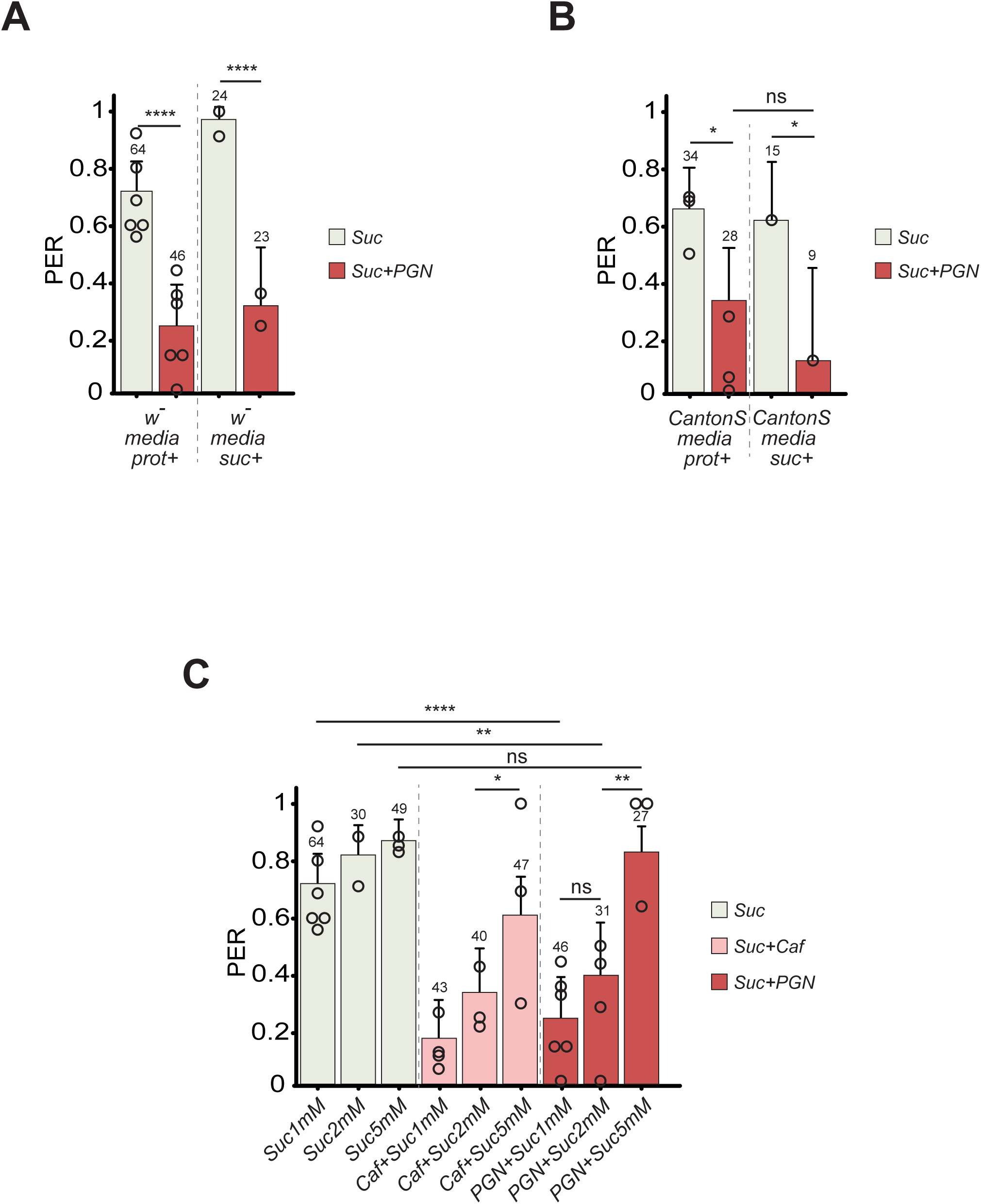
(**A**) and (**B**), raising medium has no effect on PGN-induced PER suppression. PER index of w-(**A**) and CantonS (**B**) flies, raised on sucrose (suc+) or amino acids (prot+) enriched media, to control solutions of sucrose 1mM and to sucrose 1mM + PGN from *E. coli* K12 at 200µg/mL. The data for w- raised on media prot+ are the same as figure 1A, and those for CantonS raised on media prot+ are the same as figure 1C. (**C**) Sucrose concentrations above 5mM mask the aversive effect of PGN and caffeine. PER index of w- flies to increasing concentrations of sucrose, caffeine10mM + increasing concentration of sucrose and PGN from *E. coli* K12 at 200µg/mL + increasing concentrations of sucrose. For (**A**), (**B**) and (**C**) PER index is calculated as the percentage of flies tested that responded with a PER to the stimulation ± 95% CI. The number of tested flies (n) is indicated on top of each bar. For each condition, at least 3 groups with a minimum of 10 flies per group were used. ns indicates p>0.05, * indicates p<0.05, ** indicates p<0.01, *** indicates p<0.001, **** indicates p<0.0001 Fisher Exact Test. Further details can be found in the detailed lines, conditions and, statistics for the figure section.

**Fig. S2.**
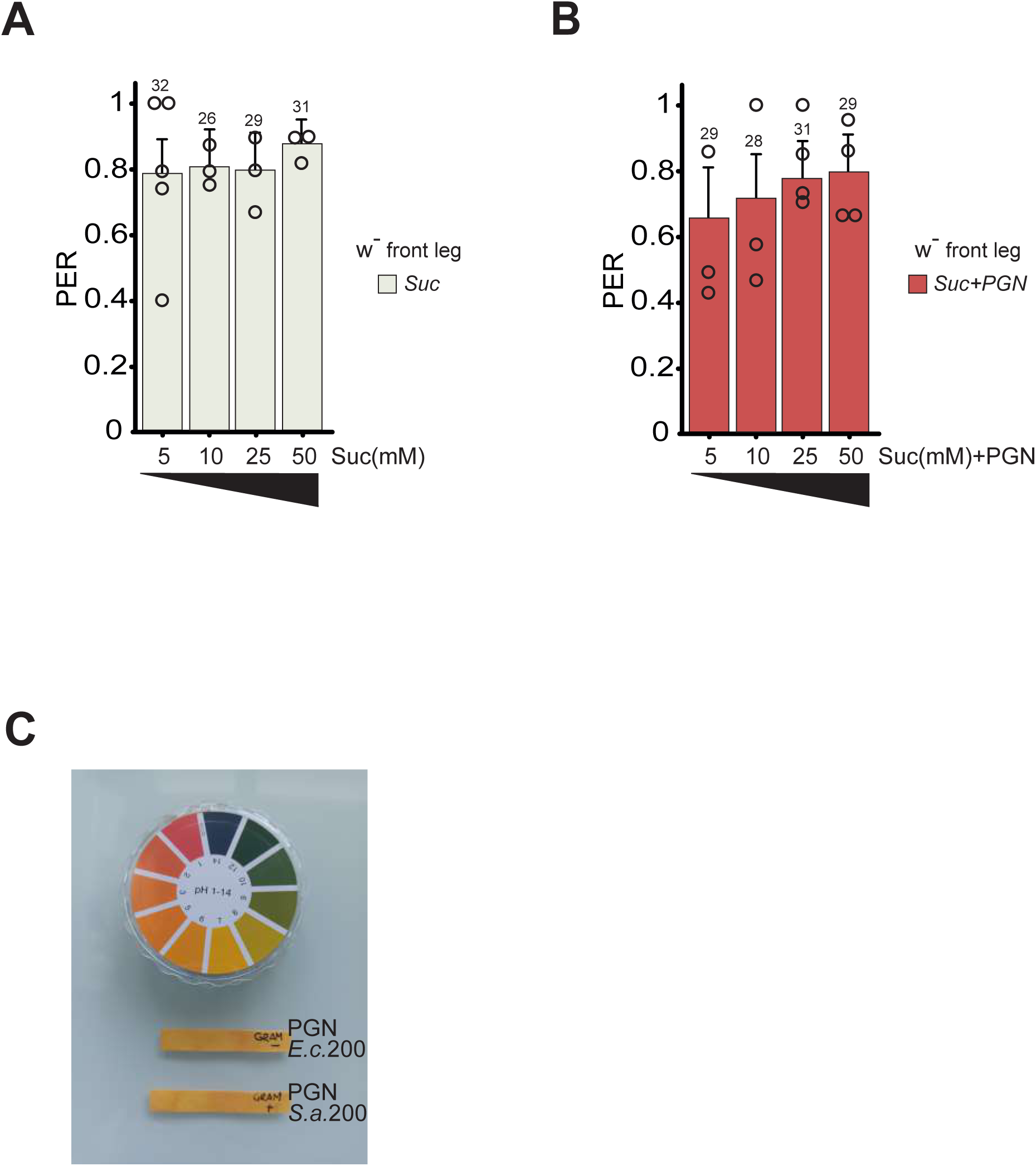
(**A**) PER index of w- flies upon stimulation of the front leg with increasing concentrations of sucrose. The numbers below the x-axis correspond to the sucrose final concentrations in mM. As 1mM sucrose does not trigger a PER robust and reproducible enough to qualify the animal as able to respond, this concentration was not used. (**B**) PER index of w- flies upon stimulation of the front leg with PGN from *E. coli* K12 at 200µg/mL + increasing concentrations of sucrose. The numbers below the x-axis correspond to the sucrose final concentrations in mM, once is mixed with PGN. (**C**) PGN solutions from *E. coli* K12 (Ec) and *S. aureus* (SA) at 200µg/mL are neutral. The pH of the two solutions was measured with pH test strips. For (**A**) and (**B**) PER index is calculated as the percentage of flies tested that responded with a PER to the stimulation ± 95% CI. The number of tested flies (n) is indicated on top of each bar. For each condition, at least 3 groups with a minimum of 10 flies per group were used. ns indicates p>0.05, * indicates p<0.05, ** indicates p<0.01, *** indicates p<0.001, **** indicates p<0.0001 Fisher Exact Test. Further details can be found in the detailed lines, conditions and, statistics for the figure section.

**Fig. S3.**
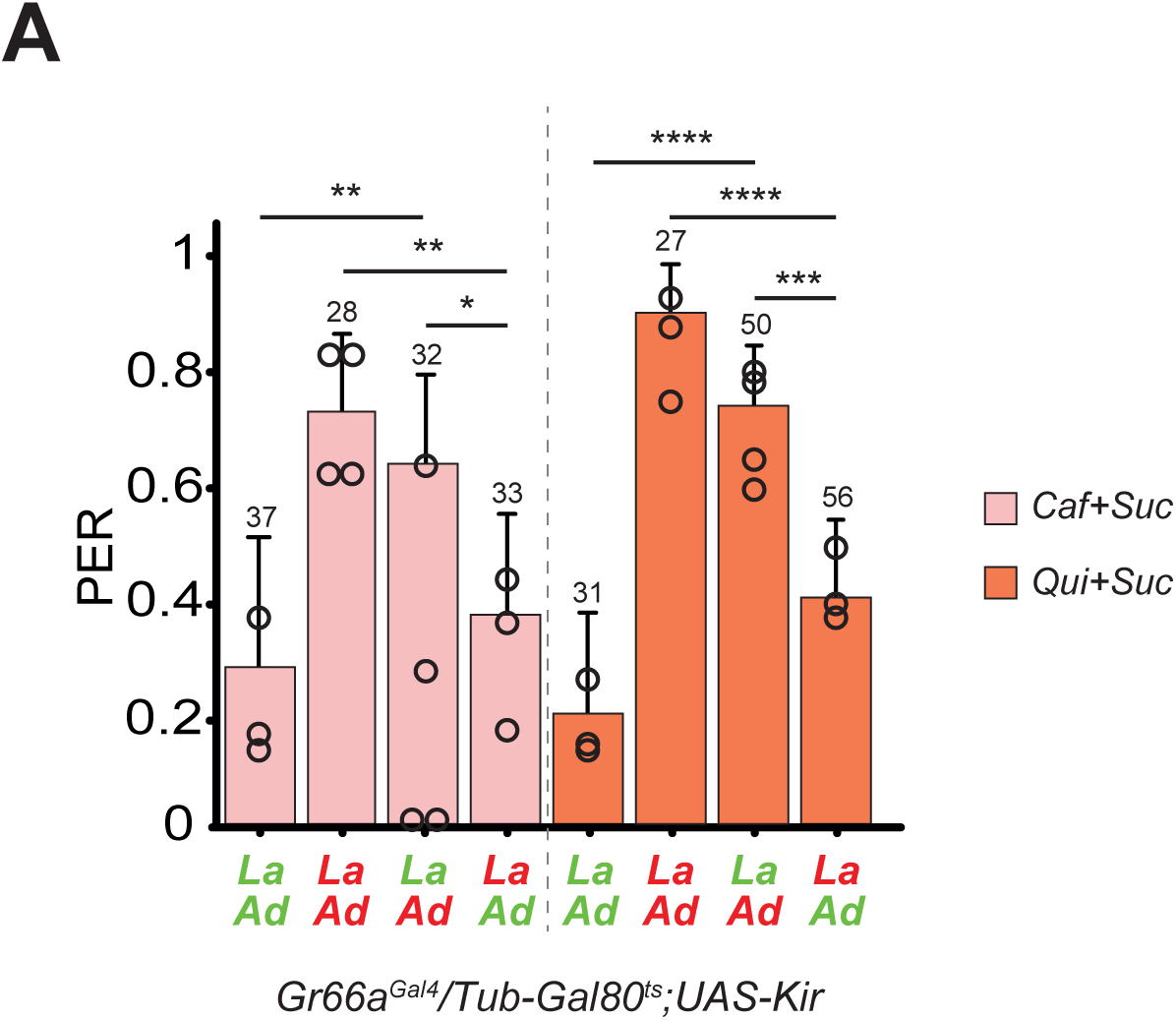
(**A**) Perception by the adult flies of sucrose 1mM + quinine 10mM and sucrose 1mM + caffeine 10mM required Gr66a+ neurons to be functional in the adult but not in larvae. The ubiquitously expressed Tub-Gal80^ts^, that inhibits the activity of Gal4, is temperature sensitive: it’s active at 18**°**C and inactivated at 29**°**C, allowing the expression of UASKir2.1 and the consequent impairment of Gr66a+ or ppk23+ neuron activity. The PER index is calculated as the percentage of flies tested that responded with a PER to the stimulation ± 95% CI. The number of tested flies (n) is indicated on top of each bar. ns indicates p>0.05, * indicates p<0.05, ** indicates p<0.01, *** indicates p<0.001, **** indicates p<0.0001 Fisher Exact Test. Further details can be found in the detailed lines, conditions and, statistics for the figure section

**Fig. S4.**
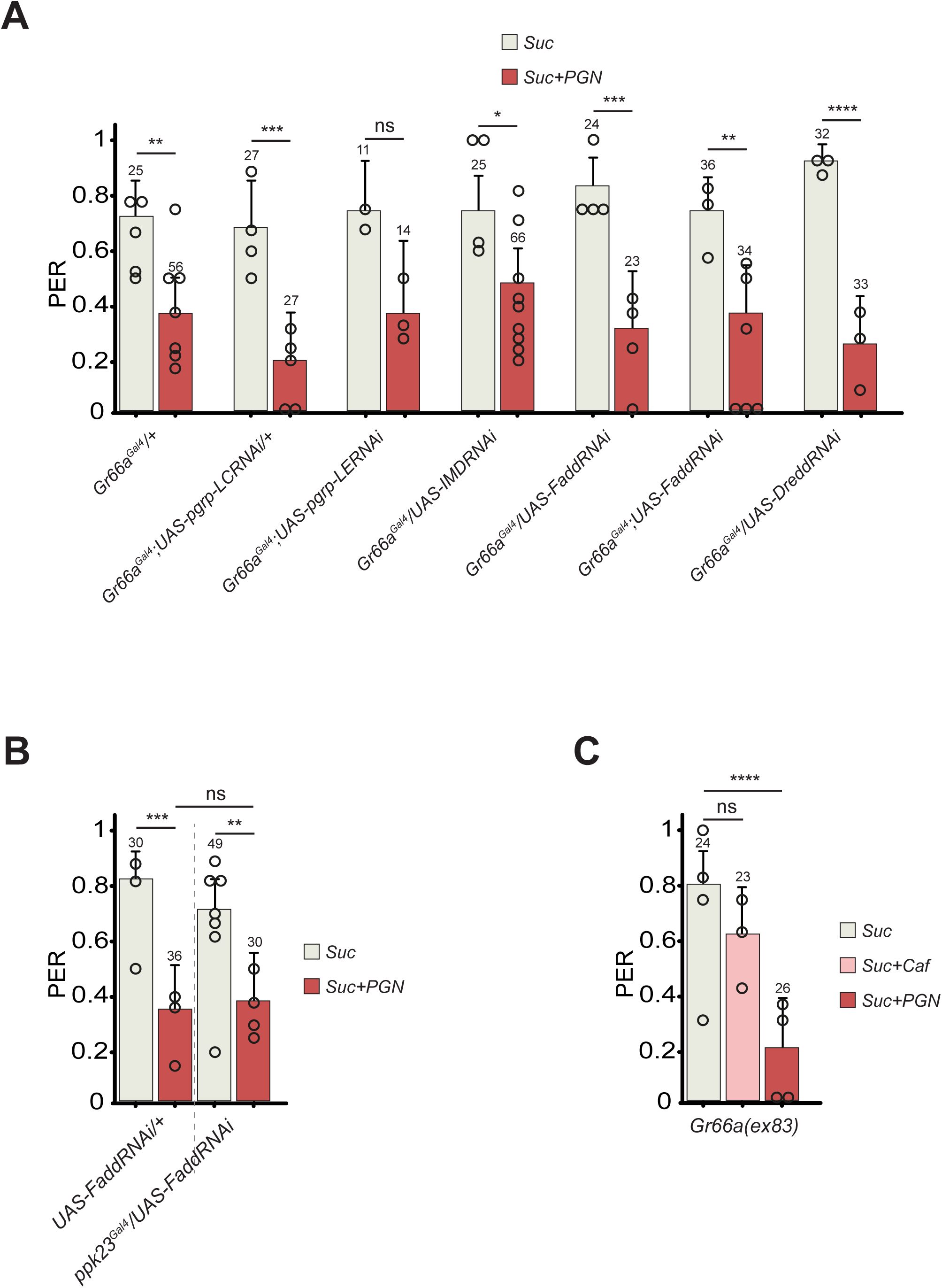
(**A**) and (**B**) RNAi-mediated inactivation of different elements of the IMD pathway has no effect on the PGN induced PER suppression. (**A**) PER index of flies in which different elements of the IMD pathway are inactivated via RNAi in Gr66a+ neurons, to control solutions of sucrose and sucrose 1mM + PGN from *E. coli* K12 at 200µg/mL. (**B**) PER index of flies upon RNAi-mediated Fadd (UAS-Fadd RNAi) inactivation in ppk23+ cells, to control solutions of sucrose and sucrose + PGN from *E. coli* K12 at 200µg/mL. (**C**) The Gr66a receptor is not involved in PGN induced PER suppression, Gr66a mutants lose their aversion to caffeine but retain their aversion to PGN. PER index of fly mutant for the Gr66a receptor to control solutions of sucrose 1mM and sucrose 1mM + caffeine and to sucrose + PGN from *E. coli* K12 at 200µg/mL. For (**A**), (**B**) and (**C**) PER index is calculated as the percentage of flies tested that responded with a PER to the stimulation ± 95% CI. The number of tested flies (n) is indicated on top of each bar. For each condition, at least 3 groups with a minimum of 10 flies per group were used. ns indicates p>0.05, * indicates p<0.05, ** indicates p<0.01, *** indicates p<0.001, **** indicates p<0.0001 Fisher Exact Test. Further details can be found in the detailed lines, conditions and, statistics for the figure section.

**Fig. S5.**
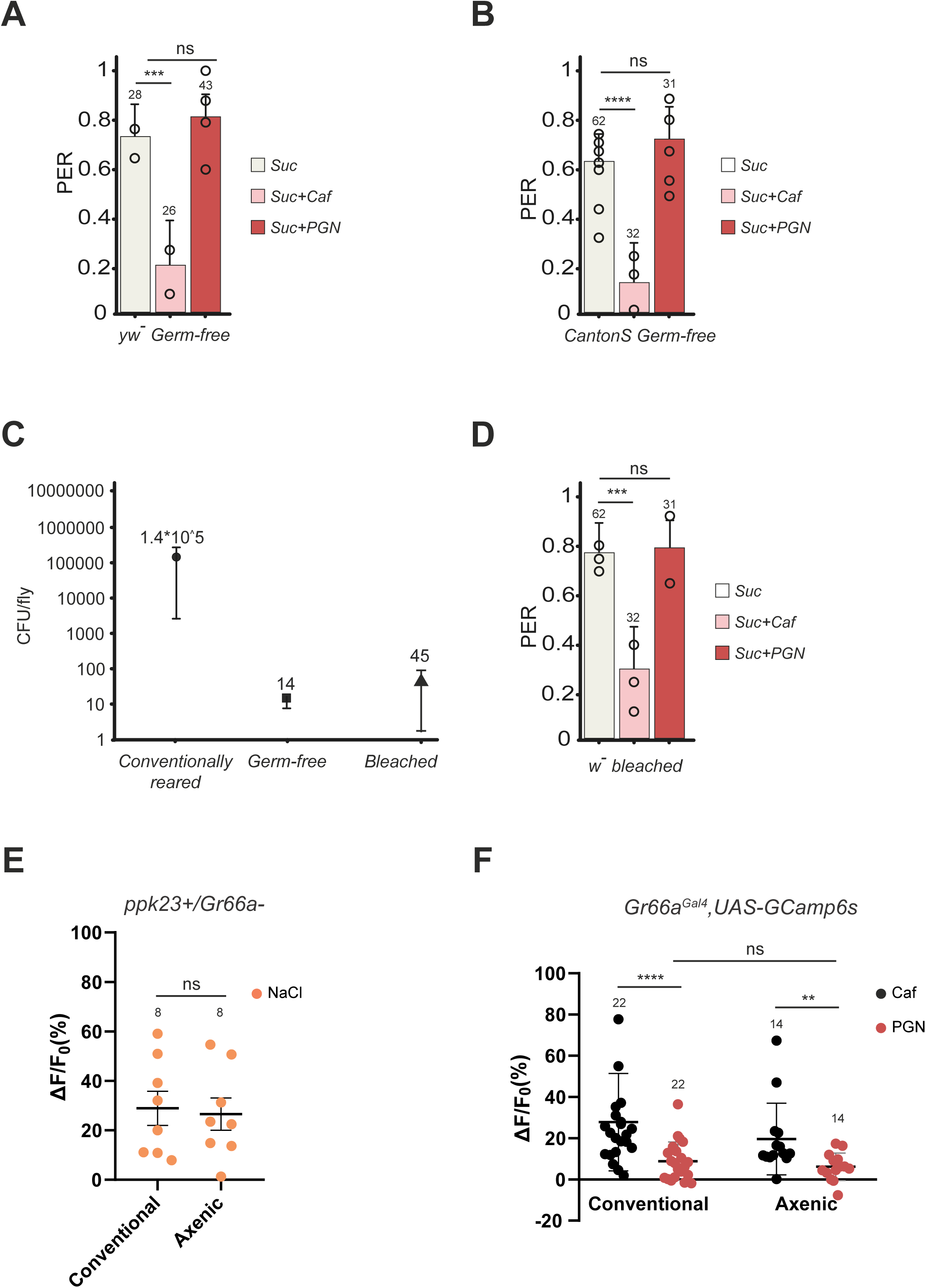
(**A)** and (**B**) Loss of aversion to PGN in axenic conditions is independent of genetic background. PER index of yw- (**A**) and CantonS (**B**) axenic flies to control solutions of sucrose 1mM and sucrose 1mM +caffeine 10mM and to sucrose 1mM + PGN from *E. coli* K12 at 200µg/mL. (**C**) and (**D**) the antibiotic treatment has no effect on the fly taste for PGN. (**C**) Bacterial load (CFU/fly) of w- flies reared under conventional conditions, germ-free with antibiotic treatment or sterilized by embryo bleaching. The data for conventionally raised and germ-free flies are the same as fig. 5A. (**D**) Flies emerged from bleached embryos retain the PGN induced PER suppression, PER index of w- flies sterilized by embryos bleaching to control solutions of sucrose 1mM and sucrose 1mM +caffeine 10mM and to sucrose 1mM + PGN from *E. coli* K12 at 200µg/mL. (**E**) Real-time calcium imaging using the calcium indicator GCaMP6s to reflect the *in vivo* neuronal activity of Gr66a+ neurons (Gr66a^Gal4^, UAS-GCaMP6s) in adult brains of conventionally raised and germ-free flies whose proboscis has been stimulated with PGN. (**E**) Averaged fluorescence intensity of positive peaks ± SEM for conventionally raised (n= 22 flies) and germ-free flies (n= 14 flies) in response to caffeine 10mM and peptidoglycan from *E. coli* K12 at 200µg/mL. (**F**) Averaged fluorescence intensity of positive peaks ± SEM for conventionally raised (n= 8) and germ-free flies (n=8) in response to Sodium chloride 250mM. For (**A**), (**B**) and (**D**) PER index is calculated as the percentage of flies tested that responded with a PER to the stimulation ± 95% CI. The number of tested flies (n) is indicated on top of each bar. For each condition, at least 3 groups with a minimum of 10 flies per group were used. ns indicates p>0.05, * indicates p<0.05, ** indicates p<0.01, *** indicates p<0.001, **** indicates p<0.0001 Fisher Exact Test. In (**E**) and (**F**), ns indicates p>0.05, ** indicates p<0.001, **** indicates p<0.00001 non-parametric t-test, Mann-Whitney test. Further details can be found in the detailed lines, conditions and, statistics for the figure section.

**Fig. S6.**
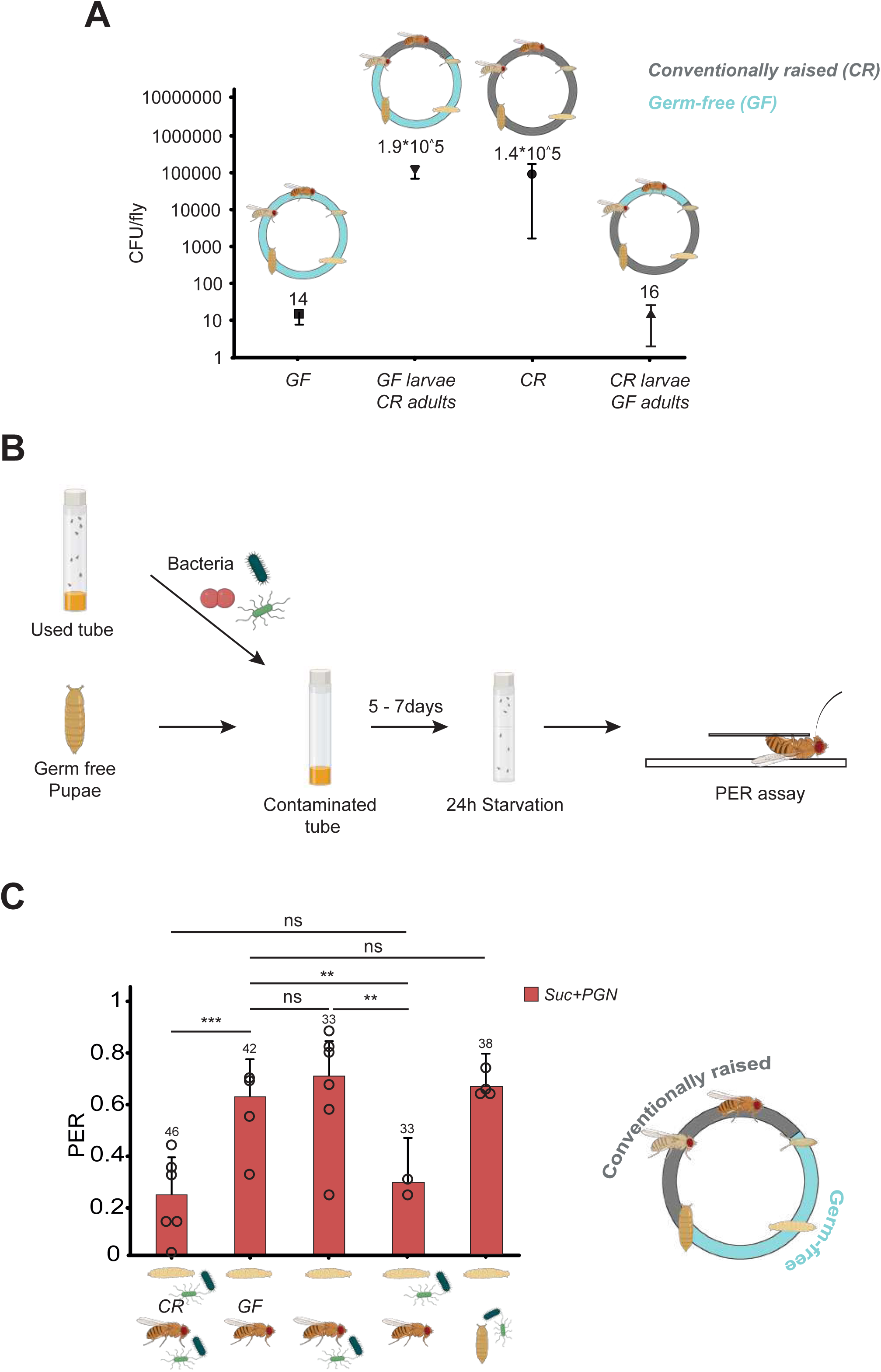
(**A**) Bacterial load (CFU/fly) of w- flies reared under conventional conditions, germ-free or shifted between the two upon pupation. The data for conventionally raised and germ-free flies are the same as fig. 5A. (**B**) Graphical representation of the protocol for pupae contamination. yw- flies are reared under conventional conditions, after a few days the fly tube is filled with Luria Bertani’s liquid medium and used as an inoculum to start a bacterial culture. The culture is then diluted to OD1 and used to soak filter paper placed over the fly culture medium in a new tube. The plug containing the germ-free pupae is dipped into the OD1 bacterial solution and used to close the previously prepared contaminated tube. (**C**) Contamination from the pupal stage is not sufficient to elicit PGN induced PER suppression. PER index of flies, that developed as germ-free larvae and were contaminated at the pupal stage, to sucrose 1mM + PGN from *E. coli* K12 at 200µg/mL. The data for flies raised in conventional conditions, germ free and for those switched between the two are the same as in Fig.6B. For (**C**) PER index is calculated as the percentage of flies tested that responded with a PER to the stimulation ± 95% CI. The number of tested flies (n) is indicated on top of each bar. For each condition, at least 3 groups with a minimum of 10 flies per group were used. ns indicates p>0.05, * indicates p<0.05, ** indicates p<0.01, *** indicates p<0.001, **** indicates p<0.0001 Fisher Exact Test. Further details can be found in the detailed lines, conditions and, statistics for the figure section;

**Fig. S7.**
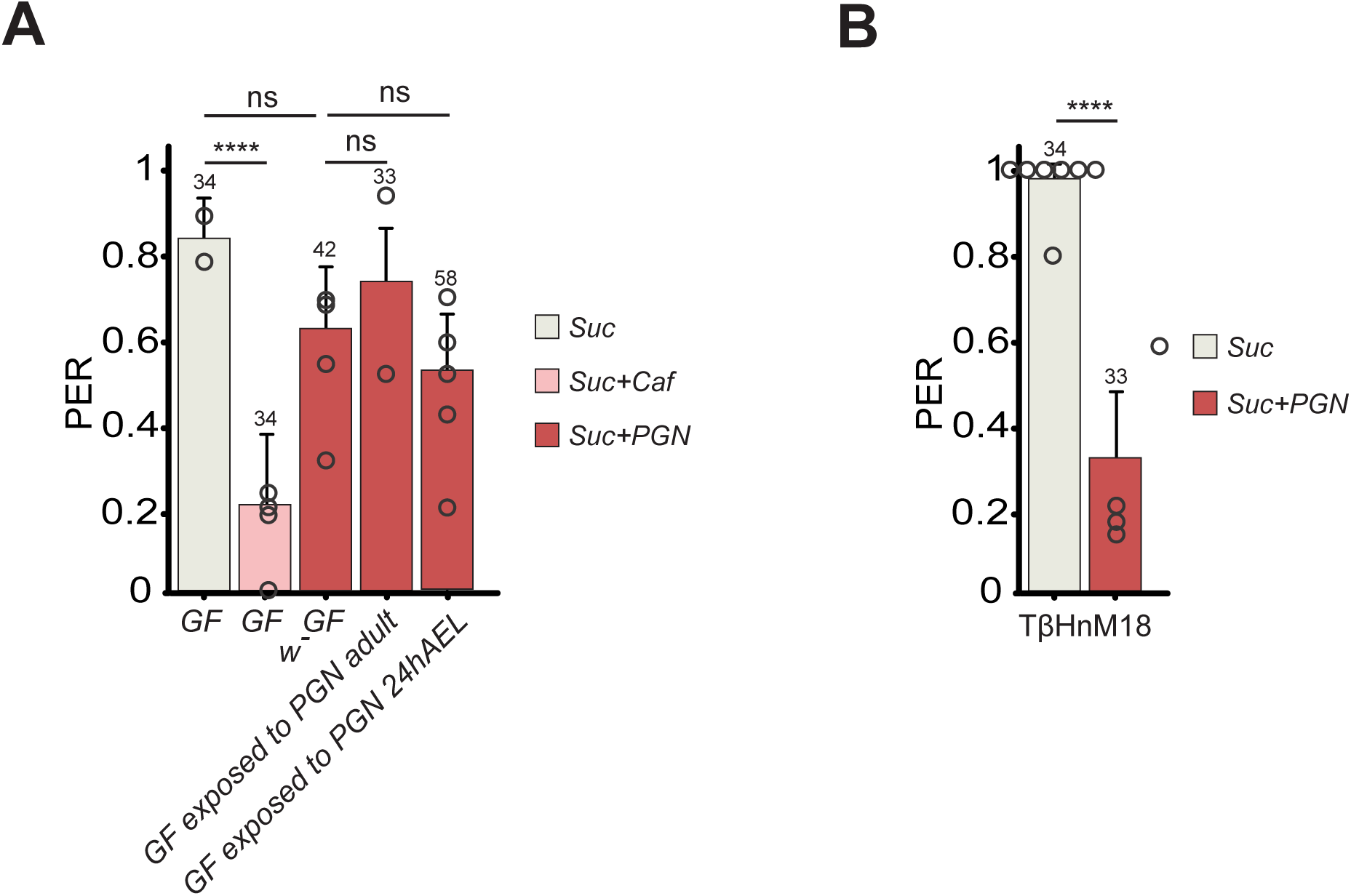
Exposing germ-free larvae and adults to PGN-containing media it is not sufficient to trigger PGN induced PER suppression. (**A**) PER index of germ-free flies exposed to PGN as a larva or as adult, to sucrose 1mM + PGN from *E. coli* K12 at 200µg/mL. The data for germ-free flies (GF) are the same as in Fig.5B. (**B**) Octopamine is not involved in PGN induced PER suppression. PER index of TßH mutant flies unable to synthetize octopamine, to control sucrose solution and to sucrose 1mM + PGN from *E. coli* K12 at 200µg/mL. For (**A**) and (**B**) PER index is calculated as the percentage of flies tested that responded with a PER to the stimulation ± 95% CI. The number of tested flies (n) is indicated on top of each bar. For each condition, at least 3 groups with a minimum of 10 flies per group were used. ns indicates p>0.05, * indicates p<0.05, ** indicates p<0.01, *** indicates p<0.001, **** indicates p<0.0001 Fisher Exact Test. Further details can be found in the detailed lines, conditions and, statistics for the figure section.

**Fig. S8.**
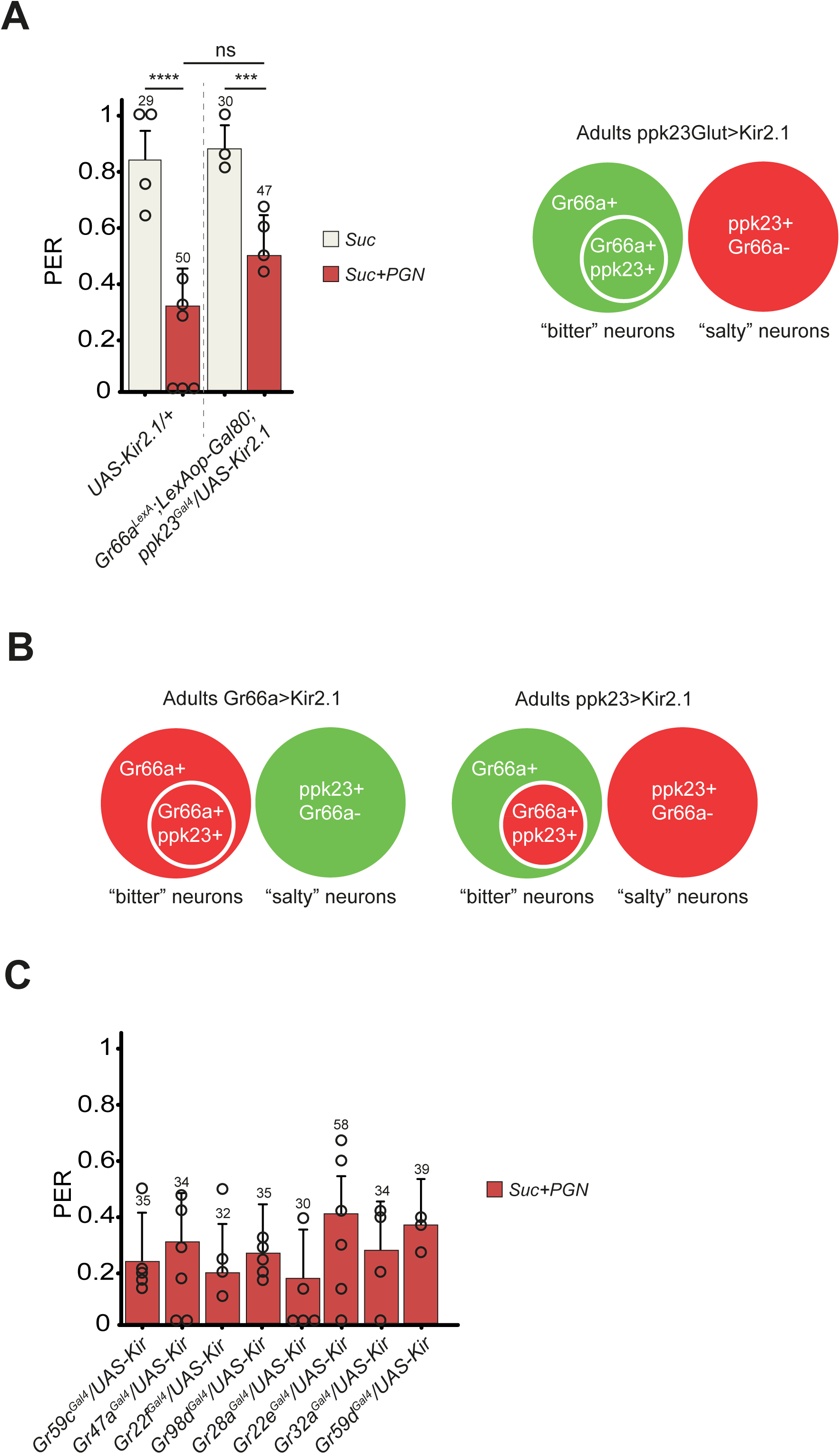
(**A**) Flies in which Gr66a-/ppk23+ cells are silenced remains able to sense PGN. PER index of flies to control solutions of sucrose and sucrose 1mM + PGN from *E. coli* K12 at 200µg/mL. The expression of LexAop-Gal80 antagonizes the activity of Gal4, thus preventing the expression of Kir2.1 in Gr66a+ neurons. The scheme represents in red the neurons inactivated by Kir2.1 and in green the active ones. ’ppk23Glut’ indicates *ppk23*^Gal4^ with *Gr66a^LexA^* and *LexAop- Gal80*; and ‘>’ denotes Gal4 driving *UAS-Kir2.1* with temporal restriction by *tub-Gal80^ts^*. (**B**) The scheme represents the result of driving the expression of Kir2.1 in Gr66a+ or ppk23+ neurons, in red the neurons inactivated by Kir2.1 and in green the active ones. ‘>’ denotes Gal4 driving *UAS-Kir2.1*. (**C**) Silencing of smaller subgroup of the Gr66a+ neurons population has no effect on the PGN-induced PER suppression. PER index of flies in which part of Gr66a+ neurons are inactivated via different Gr drivers guiding the expression of Kir2.1, to solutions of sucrose 1mM + PGN from *E. coli* K12 at 200µg/mL. For (**A**) and (**C**) PER index is calculated as the percentage of flies tested that responded with a PER to the stimulation ± 95% CI. The number of tested flies (n) is indicated on top of each bar. For each condition, at least 3 groups with a minimum of 10 flies per group were used. ns indicates p>0.05, * indicates p<0.05, ** indicates p<0.01, *** indicates p<0.001, **** indicates p<0.0001 Fisher Exact Test. Further details can be found in the detailed lines, conditions and, statistics for the figure section.

## Material and methods

### Fly husbandry

Flies were grown at 25°C on a yeast/cornmeal medium in 12h/12h light/dark cycle-controlled incubators. For 1 L of food, 8.2 g of agar (VWR, cat. #20768.361), 80 g of cornmeal flour (Westhove, Farigel maize H1) and 80 g of yeast extract (VWR, cat. #24979.413) were cooked for 10 min in boiling water. 5.2 g of Methylparaben sodium salt (MERCK, cat. #106756) and 4 mL of 99% propionic acid (CARLOERBA, cat. #409553) were added when the food had cooled down.

Schnorrer food recipe. For 1 L of food, 11 g of agar (VWR, cat. #20768.361), 80 g of cornmeal flour (Westhove, Farigel maize H1), 20 g of yeast extract (VWR, cat. #24979.413) and 30g of Sucrose were cooked for 10 min in boiling water. 2.5g of Moldex and 5 mL of 99% propionic acid (CARLOERBA, cat. #409553) were added when the food had cooled down.

### PER assay

All flies used for the test were females between 5 and 7 days old. Unless experimental conditions require it, the flies are kept and staged at 25°C to avoid any temperature changes once they are put into starvation. The day before, the tested flies are starved in an empty tube with water-soaked plug for 24h at 25°C.

Eighteen flies are tested in one assay, 6 flies are mounted on one slide and in pairs under each coverslip. To prepare the slide, three pieces of double-sided tape are regularly spaced on a slide. Two spacers are created on the sides of each piece of tape by shaping two thin cylinders of UHT paste. To avoid the use of carbon dioxide flies are anesthetized on ice. Under the microscope, two flies are stuck on their backs, side by side, on same piece of tape so that their wings adhere to the tape. A coverslip is then placed on top of the two flies and pressed onto the UHT paste, blocking their front legs and immobilizing them.

Once all slides are prepared, they are transferred to a humid chamber and kept at 25°C for 1.5 hours to allow the flies to recover before the assay.

Flies are tested in pairs, the test is carried out until completion on a pair of flies (under the same coverslip), and then move on to the next pair.

Before the test, water is given to each pair of flies to ensure that the flies are not thirsty and do not respond with a PER to the water in which the solutions are prepared. Stimulation with the test solution is always preceded and followed by a control stimulation with a sweet solution, to assess the fly’s condition and its suitability for the test. During the test small strips of filter paper are soaked in the test solution and used to contact the fly’s labellum (three consecutive times per control and test phase). Contact with the fly’s proboscis should be as gentle as possible. Ideally the head should not move. A stronger touch may prevent the fly from responding to subsequent stimulation. Based on the protocol needed the test is done following the sequence and the timing in the table below.

**Tab. 1.** Aversion protocol

|  |  |  |  |  |  |  |  |  |  |  |
| --- | --- | --- | --- | --- | --- | --- | --- | --- | --- | --- |
| Water To satiety | Wait 1 min | CON-TROL Sucrose 1mM Repeated 3 times | Wait 1 min | Water one time | Wait 1 min | TEST Repeated 3 times | Wait 1 min | Water one time | Wait 1 min | CON-TROL Sucrose 1mM Repeated 3 times |

All solutions to be tested are prepared the test day and stored at room temperature. In the aversion protocol, the control stimulation is performed with 1mM sucrose (D(+)-sucrose ≥99.5 %, p.a. Carl Roth GmbH + Co. KG). This concentration is sufficient to elicit a PER but is not so high as to influence the response to the subsequent test stimulation. After each control or test stimulation, a water-soaked strip is used to tap the proboscis and clean it.

The response of the fly to each stimulation is recorded and averaged. Flies that respond positively (PER) to at least one of the control stimulations are considered for further analysis, the others are discarded. The PER index is calculated as the percentage of flies tested that responded with a PER to the TEST stimulation and represented as ± 95% CI.

### *In vivo* calcium imaging

*In vivo* calcium imaging experiments were performed on 5-7 day-old starved mated females. Animals were raised on conventional media with males at 25°C. Flies were starved for 20-24 h in a tube containing a filter soaked in water prior any experiments. Flies of the appropriate genotype were anesthetized on ice for 1 h. Female flies were suspended by the neck on a plexiglass block (2 x 2 x 2.5 cm), with the proboscis facing the center of the block. Flies were immobilized using an insect pin (0.1 mm diameter) placed on the neck. The ends of the pin were fixed on the block with beeswax (Deiberit 502, Siladent, 1345 209212). The head was then glued on the block with a drop of rosin (Gum rosin, Sigma-Aldrich 1346 -60895-, dissolved in ethanol at 70 %) to avoid any movements. The anterior part of the head was thus oriented towards the objective of the microscope. Flies were placed in a humidified box for 1 h to allow the rosin to harden without damaging the living tissues. A plastic coverslip with a hole corresponding to the width of the space between the two eyes was placed on top of the head and fixed on the block with beeswax. The plastic coverslip was sealed on the cuticle with two-component silicon (Kwik-Sil, World Precision Instruments) leaving the proboscis exposed to the air. Ringer’s saline (130 mM NaCl, 5 mM KCl, 2 mM MgCl2, 2 mM CaCl2, 36 mM saccharose, 5 mM HEPES, pH 7.3) was placed on the head. The antenna area, air sacs, and the fat body were removed. The gut was cut without damaging the brain and taste nerves to allow visual access to the anterior ventral part of the sub-esophageal zone (SEZ). The exposed brain was rinsed twice with Ringer’s saline. GCaMP6s fluorescence was viewed with a Leica DM600B microscope under a 40x water objective. GCaMP6s was excited using a Lumencor diode light source at 482 nm ± 25. Emitted light was collected through a 505-530 nm band-pass filter. Images were collected every 500ms using a Hamamatsu/HPF-ORCA Flash 4.0 camera and processed using Leica MM AF 2.2.9. Stimulation was performed by applying 140 µL of tastant solution diluted in water on the proboscis. A minimum of 2 independent experiments with a total n for each condition ranging from 7 to 9 were performed. Each experiment consisted of a recording of 70-100 images before stimulation and 160 images after stimulation. Data were analyzed as previously described (ref) by using FIJI (https://fiji.sc/).

### Bacterial strains and maintenance

For this study, the following bacterial strains were used: *L. brevis* (Leulier’s Lab) and *L. plantarum* (Gallet’s lab). All the strains were grown in 49mL MRS liquid media (MRS Broth Fluka analytical 69966) static at 37 °C in sealed 50mL tubes for anaerobic conditions. To concentrate bacteria and reach the requested OD, 250 mL overnight cultures were centrifuged 15 min at 2250 rcf, OD at 600nm was measured and bacteria were diluted in MRS medium to the desired concentration.

### Flies bacteria load

The flies are anesthetized on ice, ten females for each experimental condition are pulled in 1.5 mL microcentrifuge tubes. Under sterile conditions, 600 µL of Luria-Bertani liquid media (LB) is added to each tube and a sterile pestle is used to homogenize the flies. The fly homogenate is then serially diluted in LB medium and plated in triplicate on LB plates. Plates are kept overnight at 37°C to facilitate bacterial growth. After two or three days of growth, the average number of colonies per dilution is calculated and the bacterial count of the fly is determined as follows:

CFU/mL = (1000µL/volume plated) *average n of colonies

CFU/mL at the origin = CFU/mL*dilution factor

CFU/fly = (CFU/mL at the origin * volume in which you homogenized flies)/n of flies homogenized.

### Embryo sterilization

Three petri dishes are filled with 2.6% bleach, 70% ethanol (Ethanol 96° RPE Carlo Erba Ref 414638) and autoclaved purified distilled water respectively. The embryos are collected by filling the tube in which oviposition occurred with PBS and using a small brush to gently detach them from the flies’ food. A 40µm cell strainer is used to collect the embryos. The cell strainer with the embryos is then dipped into: bleach 2.6% for 5 minutes, ethanol 70% for 1 minute, purified water for 1 minute, ethanol 70% for 1 minute, purified water for 1 minute. The brush used to collect the embryos is sterilized in 2.6% bleach for ten minutes, rinsed and then used to transfer the sterile embryos onto the desired media.

### Larvae contamination

An oviposition of w- germ free flies is set up on an apple agar plate at 25°C for 4h. Simultaneously an oviposition of yw- flies is set up in a regular fly tube. After 4h the w- embryos are sterilized (see protocol above) and transferred on antibiotic enriched media. Once they reach the desired developmental stage w- larvae are filtered out of the antibiotic enriched media and rinsed off with PBS. They are then transferred in the tube in which yw- larvae are growing. In this way w- larvae are exposed to the same bacteria as yw- flies but starting at a defined developmental stage. Once the adults emerge from the pupae, they are distinguished on the basis of body color. yw- flies were exposed to the bacteria throughout their lives, whereas w- flies were only exposed from the moment of transfer.

### Pupae contamination

yw- flies are raised in a regular tube, after a few days the flies are removed and the tube is filled with Luria Bertani liquid media (LB) and incubated at 37°C. The day after the OD is measured and the culture is diluted to OD1. 200µL of bacterial culture are added on filter paper on top of the fly media. Germ-free fly pupae attached to the cotton plug are soaked in the bacterial suspension and this plug is used to close the previously contaminated tube. This ensures that flies are exposed to the bacteria as soon as they leave the pupal case.

### Exposure of adult flies to PGN

As soon as they emerge from the pupal case, the germ-free adult flies are transferred to a test tube containing filter paper soaked in a Sucrose1mM+PGN200µg/mL solution and placed on top of the fly medium. Flies are then flipped into tubes prepared in the same way, every day for five days and then used for the PER test.

### Exposure of larvae to PGN

An oviposition of w- germ free flies is set up on antibiotic culture medium, in parallel with another oviposition of yw- flies set up on antibiotic culture medium. Once the fly media has been softened by the yw- larvae 500µl of PGN PGN200µg/mL solution are added on top of it together with 48h old w- germ- free larvae. Once emerged from the pupal case adult flies are sorted based on body color and staged for the PER assay on antibiotic enriched fly food.

### Microbiota sequencing

Five replicates of 20 larvae were considered for 16S metagenomic analysis. DNAs were extracted using DNeasy PowerSoil kit (Qiagen) by grinding the larvae in the C1 buffer of this kit with a Precellys (VWR).

The Oxford Nanopore Technologies 16S Barcoding Kit (SQK16S-024) was used to amplify and sequence the full-length 16S ribosomal RNA. Starting with 20-25ng of extracted DNA, 45 cycles of PCR amplification were performed using New England Biolabs LongAmp Hot Start Taq 2X Master Mix and ONT 16S barcoded primers (27F and 1492R). After purification, the amplicons were quantified and qualified to be mixed equimolarly. Sequencing was performed on Mk1C (MinKNOW 21.10.8) using an R9 flongle and the run initiated with the high-accuracy base-calling model (Guppy 5.0.17). The run generated about 429000 reads of which more than 450,000 had a QC > 9. The reads were analysed with the dedicated EPI2ME 16S pipeline (v2022.01.07) using the following parameters: minimum score > Q10, minimum length 1000, maximum length 2200, minimum coverage 50%, 5 max target sequence and two different BLAST identity thresholds were tested (85 or 90%). This version used 22,162 sequences of reference coming from the microbial 16S rRNA NCBI RefSeq database.

## Notes

### Competing Interest Statement

The authors have declared no competing interest.

